# Polyphyletic origin and horizontal exchange drive the emergence of a novel resistance-breaking *Fusarium oxysporum* f. sp. *lactucae* race

**DOI:** 10.64898/2026.09.30.755575

**Authors:** R. Jordan Price, Emily van Brabant, Helen Bates, Hanna Mestdagh, Sascha Jenkins, Emma Vines, Monica Hofte, John Clarkson

## Abstract

- *Fusarium oxysporum* f. sp. *lactucae* (Fola), the cause of lettuce fusarium wilt, poses an increasing threat to global production due to the rapid emergence of resistance-breaking isolates. However, the evolutionary origins and mechanisms underpinning this diversification remain poorly understood.
- We characterised 103 Fola isolates using molecular profiling, selecting ten key isolates for long-read whole-genome sequencing and performed *in planta* virulence assays for selected isolates. This included the first assemblies for Fola2 and Fola3, recent resistance-breaking variants and a highly divergent isolate from France designated as a putative novel race, FolaX.
- Phylogenomic analysis revealed that Fola is polyphyletic, with lettuce pathogenicity acquired independently across divergent *F. oxysporum* clades. Despite this, all lineages share a conserved ‘PanFola’ effector set with near-identical sequences, indicating extensive horizontal gene transfer. Resistance-breaking Fola variants within established races were linked to copy number variations and targeted deletions, such as the loss of the *SIX8/PSE1* locus in Fola4+. Conversely, FolaX possessed a divergent core genome and a chimeric accessory genome derived from multiple Fola lineages via segmental chromosome transfer and transposon-mediated reshuffling.
- Our findings highlight the remarkable evolutionary plasticity of Fola and the need for breeding strategies that address the dynamic nature of effector repertoires.

## Introduction

Lettuce (*Lactuca sativa*) is a global crop with a market value exceeding €3.3 billion in Europe (Eurostat, 2019) and $2 billion in the USA (USDA-NASS, 2021). However, the soilborne pathogen *Fusarium oxysporum* f. sp*. lactucae* (Fola), the causal agent of fusarium wilt, poses a severe threat to production, often causing yield losses greater than 50% (Gilard *et al*., 2017a).

First reported in Japan in 1955 (Motohashi, 1960), Fola has since spread to major lettuce-producing regions worldwide, including China, the USA, India, and throughout Europe (Gilardi *et al*., 2017a). To date, four races have been identified based on unique virulence profiles on a set of differential lettuce cultivars (International Seed Federation, 2022), with identification for some races now supported by PCR assays (Pasquali et al., 2007; Gilardi et al., 2017a; Mestdagh et al., 2023). Fola race 1 (Fola1) is the most widespread, primarily affecting field grown lettuce in warmer climates, while races 2 (Fola2) and 3 (Fola3), are largely confined to Japan and Taiwan (Gilardi *et al*., 2017a). Fola race 4 (Fola4) emerged recently in Northern Europe, a region previously free of the disease and was first reported in protected (greenhouse) lettuce in the Netherlands in 2015 (Gilardi *et al*., 2017a) and Belgium (Claerbout *et al*., 2018) and has since spread to the UK (Taylor et al., 2019), Italy (Gilardi *et al*., 2019), Spain (Gálvez *et al*., 2023), and Portugal (Halgand *et al*., 2024). This posed a major challenge to the industry, as many Fola1-resistant cultivars are susceptible to Fola4, necessitating a rapid breeding response (Garibaldi et al., 2004; Scott et al., 2010; Murray *et al*., 2021). Fola4 is now moving from protected cultivation into open fields (Gilardi *et al*., 2019; Gálvez *et al*., 2023), and Fola1 is also being reported in protected crops in northern climates (Herrero et al., 2021; van Amsterdam *et al*., 2023). This suggests an imminent and complex scenario where Fola1 and Fola4 coexist, as already observed in several European countries (Gilardi *et al*., 2017b; Guerrero *et al*., 2020; Claerbout *et al*., 2023; Halgand *et al*., 2024).

Managing fusarium wilt relies heavily on host plant resistance, frequently explained by single Resistance (R) genes that recognise specific pathogen Avirulence (Avr) effectors (Cabral *et al*., 2019; Murray *et al*., 2021; Seki *et al*., 2022). Pathogens can overcome this resistance through genomic plasticity, by modifying or deleting Avr genes. The *F. oxysporum* genome is compartmentalised into conserved core chromosomes and highly variable, dispensable accessory regions (ARs) that are enriched with transposable elements (TEs) and effector genes that largely determine the host specificity of more than 100 *F. oxysporum formae speciales* (ff. spp.; Yu *et al*., 2023). The constant rearrangement and modification of effector genes on these ARs facilitates rapid evolution in *F. oxysporum* ff. spp., enabling new races to emerge and overcome host resistance with alarming speed (Ma *et al*., 2010; van Westerhoven *et al*., 2024). The evolution of *F. oxysporum* ff. spp. is facilitated by horizontal chromosome transfer (HCT) and horizontal gene transfer (HGT), which shuffle effector genes between *F. oxysporum* strains, enabling host jumps and the emergence of new races (Ma *et al*., 2010; van Dam and Rep, 2017; Henry *et al*., 2020).

In *F. oxysporum* ff. spp., a key group of these effectors are the Secreted in Xylem (Six) proteins, first characterised in *F. oxysporum* f. sp. *lycopersici* (Fol; Houterman *et al*., 2007). Homologues of the 15 Fol *SIX* genes have since been found in varying complements across different ff.spp (Jangir *et al*., 2021). Several of the Fol *SIX* genes have been shown to encode avirulence proteins, and gene knockouts for *SIX1* (Avr3), *SIX4* (Avr1), *SIX3* (Avr2), *SIX5*, and *SIX6* in Fol have demonstrated the role of Six proteins in virulence (Rep *et al*., 2004; Houterman et al., 2008; Houterman *et al*., 2009; Gawehns *et al*., 2014; Ma *et al*., 2015). Moreover, loss of, or mutation in the Fol Avr genes *SIX4* (Avr1), *SIX3* (Avr2) and *SIX1* (Avr3) allow the pathogen to overcome the tomato resistance genes I, I-2 and I-3, respectively (Houterman *et al*., 2007; Takken and Rep, 2010; Inami *et al*., 2012).

Our previous research established that Fola1 and Fola4 are closely related but possess distinct effector repertoires, particularly regarding *SIX* genes (Bates *et al*., 2024). While both races contain homologues of *SIX9* and *SIX14*, Fola4 isolates also have a single copy of *SIX8* (Bates *et al*., 2024). Fola1 isolates possess multiple copies of *SIX14*, but these are all inactivated by the disruptive insertion of TEs, preventing expression during infection (Bates *et al*., 2024). Ongoing evolution in Fola is further evidenced by the recent emergence of resistance-breaking Fola1 and Fola4 variants. Californian Fola1 variants have been identified that overcome lettuce resistance with one isolate (VSP-0916) causing severe wilt on cv. Costa Rica No. 4, a romaine cultivar long utilised as a benchmark for Fola1 resistance (Nayak *et al*., 2025). Just as lettuce breeding programmes had begun to incorporate resistance to Fola4, a new variant ‘FOL4+’ (Fola4+) has been reported in Belgium that overcomes Fola4-resistant cultivars, such as cv. Vestia (Mestdagh *et al*., 2025). Fola4+ is characterised by the loss of *SIX8*, which may allow evasion of detection by Fola4 host resistance mechanisms (Mestdagh *et al*., 2025). Such adaptations highlight that Fola evolution can be a localised process driven by regional selection pressures (Mestdagh *et al*., 2025; Nayak *et al*., 2025).

This study aims to investigate the evolutionary origins and genomic architecture of Fola genetic diversity, particularly in resistance-breaking variants. We hypothesised that these adaptations are driven by plasticity in the accessory genome through effector loss or rearrangement. Characterisation of over 100 Fola isolates from the USA and Europe was performed using translation elongation factor 1ɑ (*TEF1ɑ*) sequencing, race-specific PCR assays and *SIX* gene assays, identifying distinct profiles for known races, variations within Fola1 and Fola4, and a completely different profile for a novel putative race from France, designated FolaX. These data combined with initial phenotyping information from research collaborators and carried out in this study for selected isolates, allowed us to select Fola variants for further analyses by genome sequencing. Comparative genomics was performed on 10 new Fola genomes, including Fola1 and Fola4 variants, and the first assemblies for Fola2 and Fola3. Our phylogenomic analysis reveals that Fola is polyphyletic, with lettuce pathogenicity acquired independently across the *F. oxysporum* species complex (FOSC). We also identify regions of effector deletion and HGT that drive these pathogenicity profiles, providing crucial knowledge for developing precise diagnostics and targeted breeding strategies for durable resistance.

## Materials and Methods

### Fungal isolates

A collection of 103 Fola isolates from different locations in Europe and the USA, was obtained from research collections, industry collaborators or our own isolations from host plants (Table S1). Fungal cultures were grown on potato dextrose agar (PDA) at 20°C and preserved as spore suspensions on plastic beads in potato dextrose broth (PDB) amended with glycerol (20% v/v) at -70°C.

### Molecular profiling of Fola isolates

Fola2 and Fola3 can be distinguished from Fola1, Fola4 and from each other on the basis of *TEF1ɑ* sequence (Taylor *et al*., 2016), while Fola1 and Fola4 are differentiated by their *SIX* gene complement and race-specific PCR assays (Pasquali *et al*., 2007; Fujinaga *et al*., 2014; Gilardi *et al*., 2017a; Bates *et al*., 2024). A total of 103 Fola isolates (49 USA, 37 Spain, 5 France, 5 Italy, 3 The Netherlands, 1 UK) as well as one Fola2 and two Fola3 isolates (Japan) were profiled using previously published or modified primers (Table S2). DNA was extracted from 20 mg of lyophilised mycelium using the DNeasy Plant kit (Qiagen). PCR amplification of *TEF1ɑ*, *SIX8*, *SIX9* and *SIX14* was carried out using REDTaq ReadyMix (Sigma-Aldrich) as per manufacturer’s guidelines, using previously published primers at the appropriate annealing temperature (Table S2). Amplicons were visualised using gel electrophoresis, purified using the QIAquick PCR Purification Kit (Qiagen) and sequenced by Eurofins using both forward and reverse primers to make a consensus sequence for each isolate.

### Virulence testing of Fola variants on lettuce plants

Four Fola1 variants (AJ853, AJ855, AJ932, AJ1004) and the putative new race FolaX (AJ934), together with standard Fola1 (AJ520) and Fola4 (AJ516) isolates as controls, were tested for virulence on five cultivars from the lettuce race differential set (cvs. Gisela, Costa Rica No. 4, Lomeria, Romabella and Palmos; Fig. S1A; International Seed Federation, 2024). Inoculum was prepared by mixing sterile Levington M2 compost and milled bran (300 g) in 1 litre conical flasks with 5 agar plugs from actively growing colonies and incubating at 23°C for 6 weeks, agitating twice weekly. This was used to infest a growing medium (John Innes No. 3) at 1×10^6^ cfu g^-1^ and used to fill 9 cm pots. Two-week-old lettuce seedlings of each cultivar were transplanted into the pots in a glasshouse compartment (25°C day, 18°C night, 16-hour photoperiod). A non-inoculated control treatment was also included for each lettuce cultivar. Each treatment had 9 replicates, giving a total of 360 pots (5 cultivars x 7 Fola isolates + 5 cultivars uninoculated) which were randomised across 9 blocks in the glasshouse. Plants were assessed twice per week for symptoms using an 8-point disease scale (Fig. S1B) for 6 weeks. Vascular browning was assessed at harvest, following longitudinal dissection of each lettuce tap root using a 4-point disease scale (Fig. S1C). The experiment was performed twice, from July to August 2024 and July to August 2025.

Statistical analysis of the data was performed in R v4.5.2 (R Core Team, 2025). Normality was assessed by a Shapiro-Wilk test. Final wilt score and vascular browning score distributions and plant dry weight data were analysed using Kruskal-Wallis tests, followed by Dunn’s post hoc test with Benjamini-Hochberg correction for multiple comparison of isolates for each cultivar. Wilt scores over time were analysed via Bayesian model (Bürkner *et al*., 2025), followed by estimated marginal means with Holm correction.

### Genomic DNA extraction

High molecular weight genomic DNA was extracted from 20 mg of lyophilised mycelium using a Macherey-Nagel NucleoSpin Plant II kit (Fisher Scientific) as per the manufacturer’s protocol. DNA purity was determined using a NanoDrop One spectrophotometer, quantity using a Qubit fluorometer, and integrity by an Agilent TapeStation TS4150. Samples were only selected for sequencing if they displayed 260/280 and 260/230 ratios between 1.8-2.0 and 2.0-2.2, respectively, a concentration Qubit:Nanodrop ratio between 0.8-1.0 and an average fragment length exceeding 50 kb.

### Genome sequencing

Long-read genome sequencing was performed using Oxford Nanopore Technologies (ONT) platform. Libraries were prepared from 1 µg of DNA using the Ligation Sequencing Kit (LSK114) according to the manufacturer’s protocol. Sequencing was performed on a GridION instrument with R10.4.1 flow cells, using high-accuracy base calling. Prior to genome assembly, all sequencing data were reprocessed using the super-high accuracy base calling model.

### Genome assembly

Quality of the ONT data was assessed using NanoPlot v1.30.1 (De Coster *et al*., 2018), and genome size was estimated with Jellyfish v2.3.0 (Marçais and Kingsford, 2011) and GenomeScope2 (Ranallo-Benavidez et al., 2020). Adapters were trimmed with Porechop v0.2.4 (Wick, 2018), followed by removing reads shorter than 1 kb or with a quality score below Q9 using Filtlong v0.2.1 (Wick, 2021). Assemblies were performed with NECAT v0.0.1 (Chen *et al*., 2021), using the estimated genome sizes and default parameters, for all isolates apart from Fola1 variant AJ853 which was assembled with Flye v2.9.5 (Kolmogorov *et al*., 2019). To correct errors, reads were aligned to the assembly using Minimap2 v2.17 (Li, 2018), informing one round of polishing with Racon v1.4.20 (Vaser *et al*., 2017) and one round with Medaka v1.5.0 (Oxford Nanopore Technologies, 2023). Assemblies were evaluated using gfastats v1.3.6 (Formenti *et al*., 2022), Merqury (Rhie *et al*., 2020), and BUSCO v5.2.2 (Simão *et al*., 2015) against the hypocreales_odb10 database. Contamination screening was performed using Kraken2 (Wood et al., 2019). Mitochondrial contigs were identified by aligning sequencing data to each assembly with LRA v1.3.7.1 (Ren and Chaisson, 2021) and calculating average coverage with Samtools v1.21 (Danecek *et al*., 2021). High coverage contigs were verified through BLASTn (Camacho *et al*., 2009) searches against the NCBI mitochondrial database. TEs and repeat sequences were annotated using EarlGrey v6.3.0 (Baril et al., 2024), with default parameters except for the “-r fungi” and “-e yes” flags. To detect core and AR contigs, a modified version of an all-vs-all whole genome alignment pipeline (van Westerhoven *et al*., 2024) was used. NUCmer v3.1 (Kurtz *et al*., 2004) was used to perform whole genome alignments between all combinations of Fola isolates and 12 non-lettuce-infecting *F. oxysporum* strains (Table S3). Alignments under 5 kb were removed and only the best match for each alignment was retained. The average number of alignments per contig were calculated and an average number of alignments cutoff of 15 was used to delineate between core and AR contigs. To confirm assignment, a principal component analysis (PCA) using the average number of alignments, GC percentage, repeat content and coverage per contig, was performed. The pairwise shared accessory content between ARs was calculated using available scripts (van Westerhoven *et al*., 2024).

### Core gene phylogenomics

Phylogenomic analysis used 3,804 complete single-copy orthologous genes to resolve relationships between Fola isolates and non-lettuce-infecting *F. oxysporum* strains (Table S3; McGowan, 2025). A maximum-likelihood phylogeny was inferred using IQ-TREE v1.6 (Nguyen *et al*., 2015), with the best-fit model selected by ModelFinder (-m MFP+MERGE; Kalyaanamoorthy *et al*., 2017) and branch support assessed with 1,000 ultrafast bootstraps (-B 1000). The final tree was rendered using iTOL v6 (Letunic and Bork, 2021).

### Effector prediction

A two-stage approach was used to generate a presence/absence profile of putative effector genes. First, the FoEC2 pipeline (van Dam *et al*., 2016; Brenes Guallar *et al*., 2022) was used for *de novo* prediction, with default parameters except for a maximum size of 300 amino acids. This effector set was cross referenced against an existing list of Fola candidate effectors (Bates *et al*., 2024) using BLASTn, and identical hits were removed to create a curated set. In the second stage, this curated set was used with the FoEC2 pipeline’s presence/absence mode (“-e” flag). Sequences for each effector present in all Fola isolates (‘PanFola’ effectors) were aligned with MAFFT, trimmed with TrimAL, and reconstructed into individual phylogenetic trees using IQ-TREE v1.6, with branch support assessed with 1,000 ultrafast bootstraps. A consensus phylogeny was generated using ASTRAL-Pro3 (Zhang et al., 2025). All trees were rendered using iTOL v6.

### Structural variation in Fola4 variant Fola4+

Structural variation was investigated by aligning Fola4 AJ592 and Fola4+ AJ993 filtered sequencing data to the Fola4 AJ592 genome using LRA v1.3.7.1. Samtools v1.21 was used to convert, sort and index the alignment files. The *SIX8/PSE1* (Paired with SIX8) locus was plotted using Samplot v1.3.0 (Belyeu *et al*., 2021), with the options “--min_event_size 30000” and “--same_yaxis_scales”.

### Splits networks

The genetic relatedness of the Fola core and AR was investigated using whole genome-based phylogenetic network reconstruction using SANS ambages (Rempel and Wittler, 2021). Each assembly was split into core and AR contigs, which were then used as inputs, with default k-mer length (31). The splits networks were visualised using SplitsTree v6 (Huson and Bryant, 2024).

### Investigation of potential HGT in FolaX

#### Comparative k-mer analysis

To identify inheritance across the FolaX genome, we compared k-mers across the Fola assemblies and the non-lettuce-infecting *F. oxysporum* genomes (Table S3). Canonical 31-mers were generated per isolate with Jellyfish v2.3.0. HGT-signal k-mers were defined as those present in FolaX AJ934 and at least one other Fola isolate but absent from the non-lettuce-infecting FOSC isolates. As the core-genome phylogeny placed FolaX as ancestral to a clade comprising Fola3 AJ810, *Fo* f. sp. *lini* 39, *Fo* f. sp. *conglutinans* Fo5176 and *Fo* f. sp. *coriandrii* 3-2, a second partition defined clade-exclusive k-mers as those present in FolaX and at least one other clade member but absent from all other Fola and FOSC isolates, distinguishing recently acquired HGT sequences from those vertically inherited. Both k-mer sets (shared-Fola and clade-exclusive) were mapped to the FolaX assembly with Bowtie2 v2.5.1 (Langmead and Salzberg, 2012) allowing zero mismatches. To give a conservative density estimate and avoid inflation by repetitive elements, Bowtie2 was run in default mode, reporting a single pseudo-random alignment for multi-mapping k-mers (MAPQ 0). Density was calculated in 10 kb sliding windows with bedtools coverage (Quinlan and Hall, 2010), and putative HGT and ancestral regions were defined as windows containing >500 shared canonical k-mers. Enrichment of predicted effectors within these regions was tested by Fisher’s exact test (bedtools fisher). To examine the mechanism of rearrangement, 94 boundaries were identified where the dominant signal transitioned between shared-Fola and ancestral sequences. Breakpoint proximity to known TEs was measured with bedtools closest against the FolaX repeat annotation, and the Kimura-80 distances of boundary TEs were compared with the genome-wide average for each TE class to assess relative element activity.

#### Proximity analysis of effectors with Starship elements

Starship transposons were identified across the 16 Fola genomes with Starfish v1.0.0 (Gluck-Thaler and Vogan, 2024). Predicted FolaX effector genes were mapped relative to Starship elements and Captain genes. An effector was classified as Starship-associated where its entire range fell within the predicted element boundaries. Proximity was further assessed using a 250 kb window from orphan Captain genes to capture effectors in possible unannotated elements.

#### Average nucleotide identity and whole genome alignments

Average Nucleotide Identity (ANI) was calculated for both core and AR across all Fola isolates with FastANI v1.34 (Jain *et al*., 2018). To pinpoint regions of high sequence identity, the FolaX assembly was aligned with Minimap2 v2.17 against a representative isolate from each race: Fola1 (AJ520), Fola2 (AJ808), Fola3 (AJ810) and Fola4 (AJ516). Alignments within the ARs were filtered to retain those exceeding the average AR ANI (>96% identity). Conserved pairwise regions containing effectors and their flanking repetitive elements were identified with a custom Python script to locate candidate effector genes (CEGs). The spatial distribution of these effectors and their alignments were visualised with pyCircos v0.3 (Mori, 2022) and PyGenomeViz v1.6.1 (Shimoyama, 2024).

#### Phylogenetic incongruence

Orthologs of the shared CEGs were identified in the non-lettuce-infecting *F. oxysporum* strains (Table S3) using BLASTn. Each CEG was aligned with MAFFT, trimmed with TrimAL and reconstructed with IQ-TREE v1.6, with branch support from 1,000 ultrafast bootstraps, and the trees were rendered in iTOL v6. A consensus phylogeny was generated with ASTRAL-Pro3 and plotted as a co-phylogeny against the core species tree using the ape v5.8.1 (Paradis *et al*., 2004) and phytools v2.5.2 (Revell, 2012) packages in R. Robinson-Foulds (RF) distances (ape v5.8.1) quantified the global symmetric difference between the two topologies. To identify candidate HGT isolates, we performed a Procrustean Approach to Co-phylogeny (PACo) and a Procrustes permutation test (protest) with the paco v0.4.2 (Hutchinson *et al*., 2017) and vegan v2.7.2 (Dixon, 2003) packages in R. Distance matrices were converted to principal coordinates under Cailliez correction to handle non-Euclidean distances, and the effector-tree coordinates were rotated and scaled onto the core species tree to maximise fit, with global significance assessed over 1,000 permutations. Individual squared residuals were then calculated per taxon to pinpoint isolates contributing to incongruence.

## Results

### Molecular characterisation of Fola isolates and selection for genome sequencing

Molecular profiling of 49 Fola isolates from the USA and 49 from Europe identified 88 as Fola1 and 7 as Fola4 (Table S1) based on *TEF1ɑ* sequence, race specific PCR assays and presence of *SIX9* and *SIX14* for Fola1 and *SIX8*, *SIX9* and *SIX14* for Fola4. *TEF1ɑ*, *SIX9* and *SIX14* sequences were identical (data not shown). Seven isolates including the published Californian Fola1 variant VSP-916 (AJ1004) matched the Fola1 profile (identical *TEF1ɑ* sequence, positive Fola1 PCR, *SIX9* present) but amplified *SIX14* only with primers within a single exon, not with standard primers spanning both exons, suggesting a modification at this locus as seen in one copy of *SIX14* in Fola1 isolate AJ520 (Bates *et al*., 2024). The identities of Fola2 (AJ808) and Fola3 (AJ809, AJ810) were confirmed, with both negative for Fola1/Fola4 PCR assays and having *TEF1ɑ* sequences matching the NCBI-deposited references for these races (Fig. S2). These isolates also resulted in amplification of *SIX8*, *SIX9* and *SIX14* with sequences identical to those found for Fola1 and Fola4. The Fola4 variant Fola4+ isolate (AJ993) was positive for the Fola4 PCR and contained *SIX9* and *SIX14* but lacked *SIX8* as reported by Mestdagh et al. (2025). Most significantly, three isolates from France (AJ933, AJ934, AJ935) had a unique molecular profile which did not match any of the known races as they i) were negative for Fola1/Fola4 specific PCRs, ii) had a *TEF1ɑ* sequence substantially different to Fola1, Fola2, Fola3 and Fola4 (Fig. S2) and iii) had a different *SIX8* sequence to that found in Fola2, Fola3 or Fola4 (Fig. S3). These isolates were considered to be a putative new race, designated FolaX.

### Virulence testing of Fola variants on lettuce plants

In both experiments testing the Fola variants across the lettuce differential cultivars, fusarium wilt symptoms developed 7–10 days post-transplant, with final wilt and vascular browning (VB) scores generally consistent across the Fola isolate/cultivar combinations (Fig. 1, Fig. S4). All isolates were highly virulent on the universally susceptible cv. Gisela, most plants had VB scores >3. The exception was FolaX AJ934, which was less virulent on cv. Gisela than the other isolates, though only significantly so in the first experiment (Fig. 1a). As expected, the standard Fola1 isolate AJ520 was also virulent on cv. Lomeria, while cvs. Costa Rica No. 4, Palmos and Romabella were resistant. In contrast, the four putative Fola1 variant isolates from the USA (Fola1v AJ853, AJ855, AJ1004) and Spain (Fola1v AJ932) were all virulent on cvs. Costa Rica No. 4 and Palmos, but not Romabella, with significantly greater VB scores (2–4) than the standard Fola1 AJ520 (0–1), indicating that they overcame these sources of Fola1 resistance (Fig. 1). This confirmed previous phenotype results for Fola1v AJ855 and AJ1004 (Table 1) and showed that AJ853 and AJ932 are also resistance-breaking variants. Significantly, the putative new race FolaX was virulent on all cultivars and so did not fit the differential profile of any known Fola race (Fig. 1).

**Figure 1.**
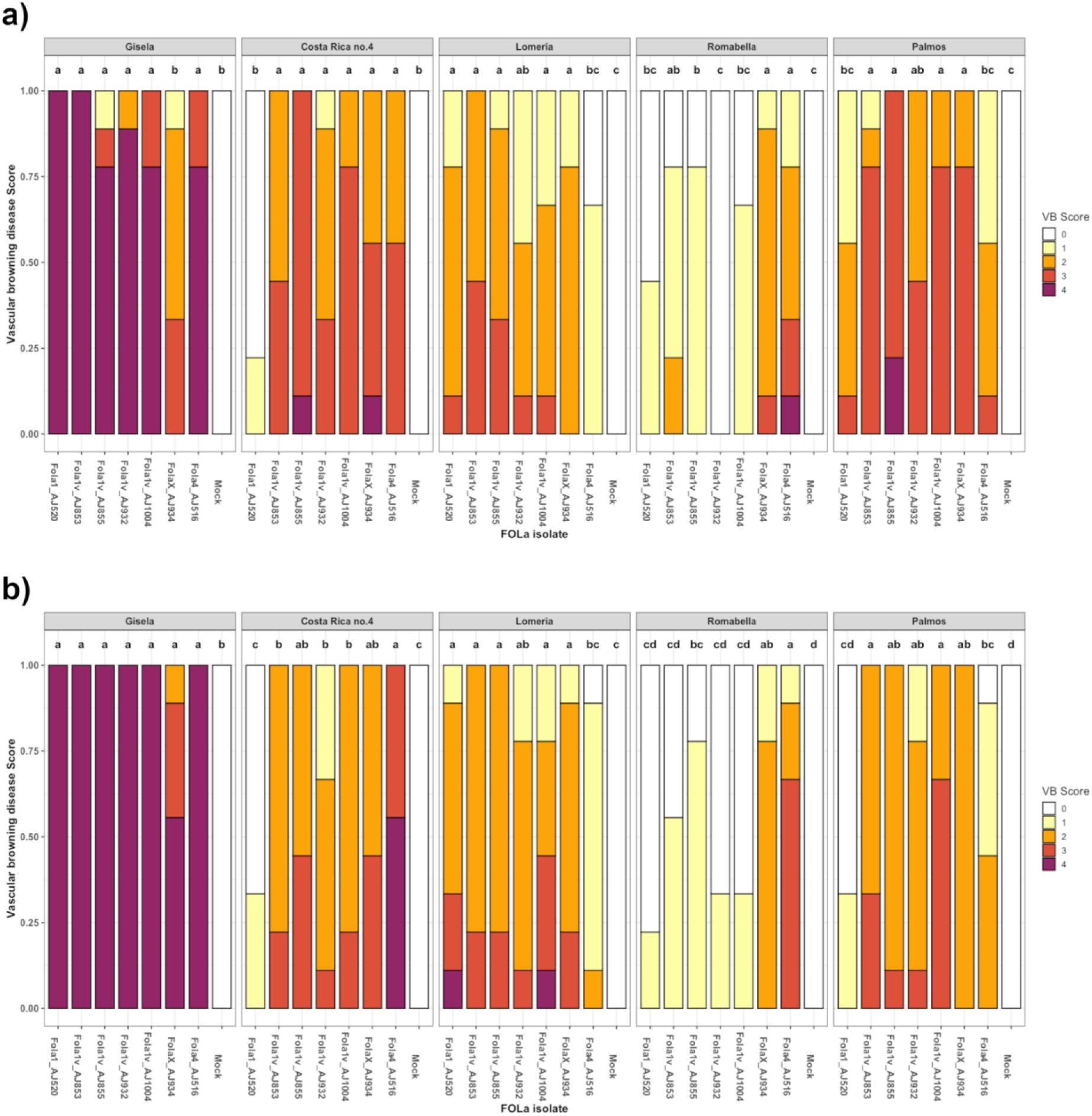
Vascular browning (VB) score distributions for Fola variant isolates on differential lettuce cultivars (Costa Rica No. 4, Gisela, Lomeria, Palmos and Romabella) for two experiments conducted in (a) 2024 and (b) 2025. Mean disease score distributions sharing a common letter are not significantly different by Kruskal-Wallis with Dunn’s post hoc test with Benjamini-Hochberg correction at the 5% significance level (“a” always denotes the most diseased plants).

**Table 1.**
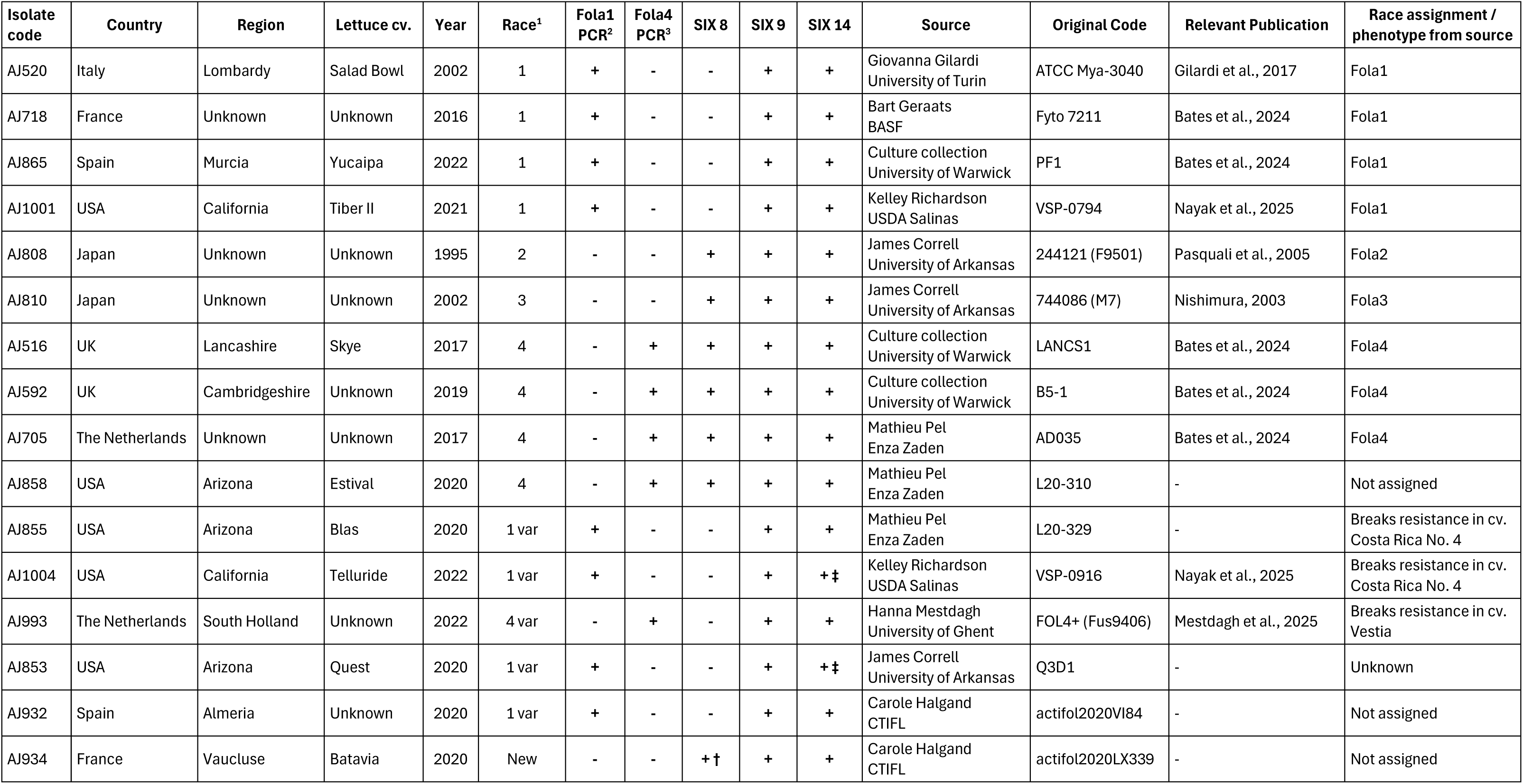
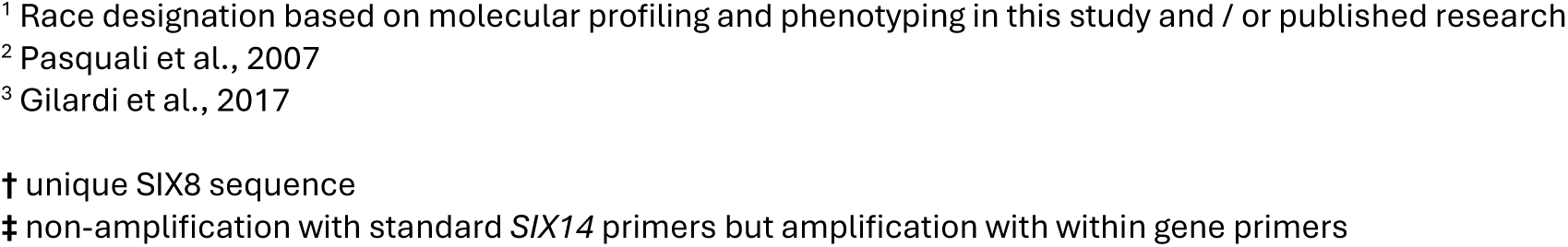
Fola isolates used in genome sequencing with details and molecular profile.

| Isolate code | Country | Region | Lettuce cv. | Year | Race<sup>1</sup> | Fola1 PCR<sup>2</sup> | Fola4 PCR<sup>3</sup> | SIX 8 | SIX 9 | SIX 14 | Source | Original Code | Relevant Publication | Race assignment / phenotype from source |
| --- | --- | --- | --- | --- | --- | --- | --- | --- | --- | --- | --- | --- | --- | --- |
| AJ520 | Italy | Lombardy | Salad Bowl | 2002 | 1 | + | - | - | + | + | Giovanna Gilardi University of Turin | ATCC Mya-3040 | Gilardi et al., 2017 | Fola1 |
| AJ718 | France | Unknown | Unknown | 2016 | 1 | + | - | - | + | + | Bart Geraats BASF | Fyto 7211 | Bates et al., 2024 | Fola1 |
| AJ865 | Spain | Murcia | Yucaipa | 2022 | 1 | + | - | - | + | + | Culture collection University of Warwick | PF1 | Bates et al., 2024 | Fola1 |
| AJ1001 | USA | California | Tiber II | 2021 | 1 | + | - | - | + | + | Kelley Richardson USDA Salinas | VSP-0794 | Nayak et al., 2025 | Fola1 |
| AJ808 | Japan | Unknown | Unknown | 1995 | 2 | - | - | + | + | + | James Correll University of Arkansas | 244121 (F9501) | Pasquali et al., 2005 | Fola2 |
| AJ810 | Japan | Unknown | Unknown | 2002 | 3 | - | - | + | + | + | James Correll University of Arkansas | 744086 (M7) | Nishimura, 2003 | Fola3 |
| AJ516 | UK | Lancashire | Skye | 2017 | 4 | - | + | + | + | + | Culture collection University of Warwick | LANCS1 | Bates et al., 2024 | Fola4 |
| AJ592 | UK | Cambridgeshire | Unknown | 2019 | 4 | - | + | + | + | + | Culture collection University of Warwick | B5-1 | Bates et al., 2024 | Fola4 |
| AJ705 | The Netherlands | Unknown | Unknown | 2017 | 4 | - | + | + | + | + | Mathieu Pel Enza Zaden | AD035 | Bates et al., 2024 | Fola4 |
| AJ858 | USA | Arizona | Estival | 2020 | 4 | - | + | + | + | + | Mathieu Pel Enza Zaden | L20-310 | - | Not assigned |
| AJ855 | USA | Arizona | Blas | 2020 | 1 var | + | - | - | + | + | Mathieu Pel Enza Zaden | L20-329 | - | Breaks resistance in cv. Costa Rica No. 4 |
| AJ1004 | USA | California | Telluride | 2022 | 1 var | + | - | - | + | + | Kelley Richardson USDA Salinas | VSP-0916 | Nayak et al., 2025 | Breaks resistance in cv. Costa Rica No. 4 |
| AJ993 | The Netherlands | South Holland | Unknown | 2022 | 4 var | - | + | - | + | + | Hanna Mestdagh University of Ghent | FOL4+ (Fus9406) | Mestdagh et al., 2025 | Breaks resistance in cv. Vestia |
| AJ853 | USA | Arizona | Quest | 2020 | 1 var | + | - | - | + | + | James Correll University of Arkansas | Q3D1 | - | Unknown |
| AJ932 | Spain | Almeria | Unknown | 2020 | 1 var | + | - | - | + | + | Carole Halgand CTIFL | actifol2020VI84 | - | Not assigned |
| AJ934 | France | Vaucluse | Batavia | 2020 | New | - | - | + | + | + | Carole Halgand CTIFL | actifol2020LX339 | - | Not assigned |

<sup>1</sup> Race designation based on molecular profiling and phenotyping in this study and / or published research
<sup>2</sup> Pasquali et al., 2007
<sup>3</sup> Gilardi et al., 2017
† unique SIX8 sequence
‡ non-amplification with standard *SIX14* primers but amplification with within gene primers

### Genome sequencing, phylogenomics & pairwise shared accessory content

Based on the molecular profiling and virulence data, a total of eight new Fola isolates were chosen for whole genome sequencing; five Fola1 isolates (including the four resistance-breaking variants; AJ853, AJ855, AJ932, AJ1004), the Fola4+ variant (AJ993), a USA isolate with typical Fola4 profile (AJ858), and one of the putative new race isolates from France (FolaX AJ934). To determine if FolaX was related to Fola2 (AJ808) or Fola3 (AJ810), we also sequenced these races for the first time (Table 1). *De novo* assembly produced highly contiguous genomes for nine isolates (N50 1.70-3.64 Mb; L50 7-12 contigs; Fig. 2a, Table S4). Limited sequencing data for AJ858 gave a more fragmented assembly (N50 0.57 Mb; L50 34 contigs), but all genomes had high quality scores (Q > 38), and high gene space completeness (complete BUSCO > 95.7%). Assemblies ranged from 59.5-66.7 Mb, with 16.0-20.7% repeat content (Fig. 2a, Table S4), comparable to our previous Fola1 and Fola4 assemblies (Bates *et al*., 2024).

**Figure 2.**
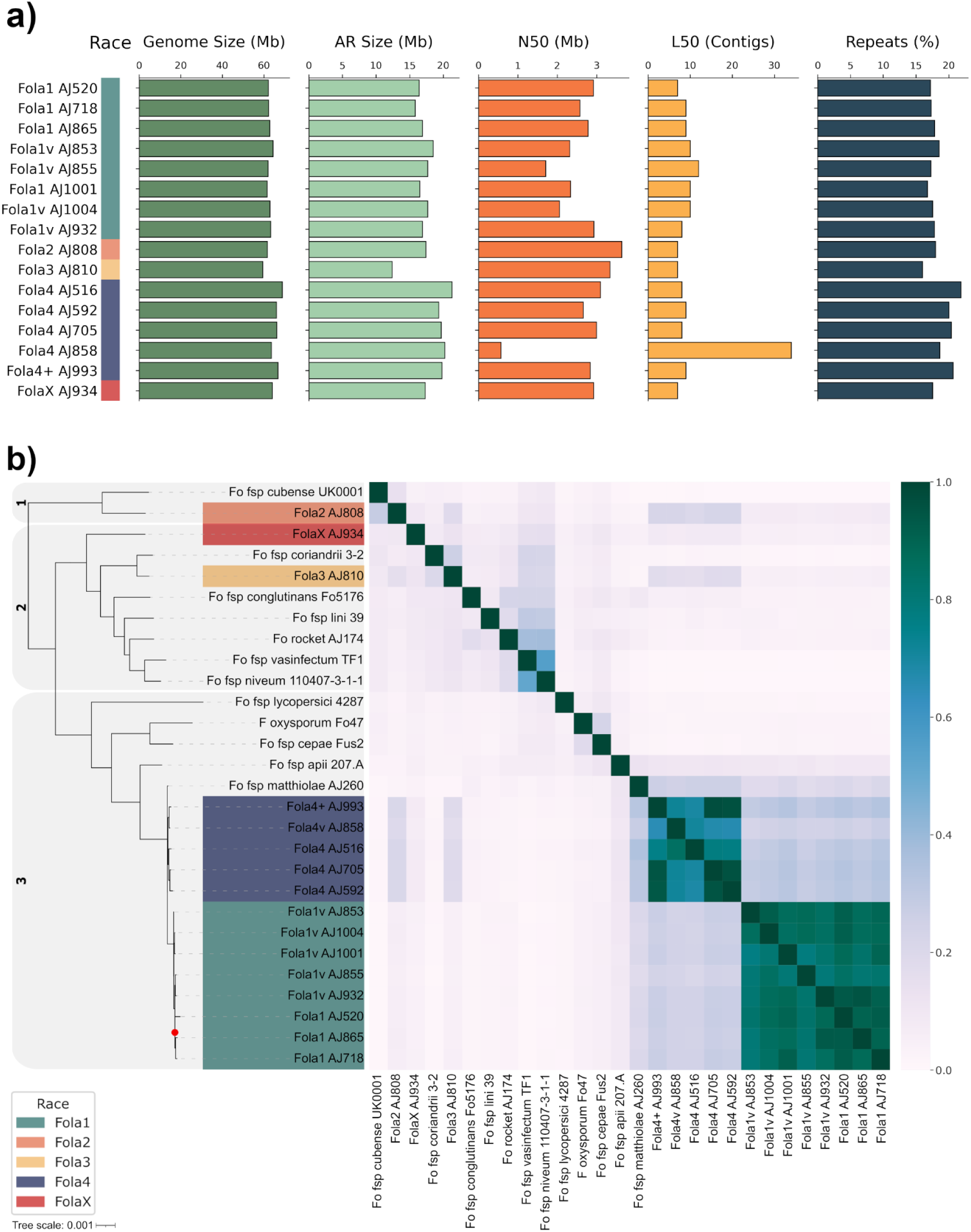
Genome assembly metrics, phylogenomic relationships and pairwise shared accessory content of *Fusarium oxysporum* f. sp. *lactucae* (Fola) isolates. (a) Bar charts detailing key assembly statistics for the sequenced Fola genomes, including total genome size (Mb), accessory region (AR) size (Mb), contig N50 (Mb), L50 (number of contigs), and repetitive element content (%). (b) Maximum-likelihood core genome phylogeny based on 3,804 single-copy orthologous genes (left), aligned with a heatmap displaying the proportion of pairwise shared accessory content between isolates (right). The Fola races are highlighted in green (Fola1), orange (Fola2), yellow (Fola3), blue (Fola4), and red (FolaX). The three FOSC clades are indicated by the shaded boxes and identified by the corresponding number. The heatmap colour scale (0.0 to 1.0) indicates the degree of AR sequence conservation, highlighting high intra-race sharedness and low inter-race sharedness.

To assess the diversity of these new isolates alongside our previous six Fola1/Fola4 assemblies and 12 high-quality FOSC genomes (contigs ≤50, BUSCO >97%), we identified 3804 single copy orthologs that were present in all isolates and constructed a maximum likelihood phylogeny (Fig. 2b). Consistent with previous molecular characterisation (Bates *et al*., 2024; Waqas *et al*., 2025), Fola isolates were polyphyletic, with Fola2 (AJ808) within FOSC Clade 1, Fola3 (AJ810) and FolaX (AJ934) in FOSC Clade 2, and Fola1 and Fola4 within FOSC Clade 3 (Fig. 2b; O’Donnell *et al*., 1998, 2009).

ARs were identified using a pangenomic approach based on all-vs-all whole-genome alignments, with contigs provisionally classified as accessory where they aligned to fewer than 15 of the other FOSC genomes (Fig. S5). This was validated by repeat density, where ARs averaged 46% repetitive content versus 8% for the core. PCA on sequence conservation, repeat density, GC content and read coverage confirmed a clear distinction between the core, accessory and mitochondrial contigs (Fig. S6). Total AR size varied considerably, from 12.4 Mb (21% of the genome) in Fola3 (AJ810) to 21 Mb (31%) in Fola4 isolate AJ516 (Fig. 2a). Pairwise comparison of shared AR content revealed high intra-race sharing (∼85% within Fola1; ∼79% within Fola4; Fig. 2b). In contrast, inter-race sharedness was low: Fola2 shared more AR content with Fola4 (22%) than Fola1 (7.5%), as did Fola3 (Fola4 17%; Fola1 5.5%; Fig. 2b). FolaX AJ934 was the most divergent, with low average AR sharedness to all races (5.3% to Fola1, 8.2% to Fola4, 9.9% to Fola2, and 12.6% to Fola3; Fig. 2b), indicating that only limited genetic material is shared across the accessory genomes of different races. Collectively, these results confirm the polyphyletic origin of Fola, with distinct lineages supported by both core genome phylogeny and low inter-race AR content.

### Candidate effectors are lost in resistance-breaking race variants

As effector gene repertoires are known to define host specificity in *F. oxysporum* ff. spp. (van Dam *et al*., 2016; Brenes Guallar *et al*., 2022), we expected effector presence/absence profiles to distinguish the Fola races. Across the 16 Fola isolates we predicted 78 MIMP-associated effector gene families, plus 24 candidates previously identified as expressed *in planta* following Fola1 and/or Fola4 infection (Bates et al., 2024). As expected, these effector genes were enriched in the ARs (Fig. S7), and the resulting repertoires clustered the isolates by race (Fig. 3). We designated the effectors expressed *in planta* in our previous study as Candidate Effector Genes (CEG01–CEG24), while those identified solely by *de novo* prediction here are termed Putative Effector Genes (PEG01–PEG78).

**Figure 3.**
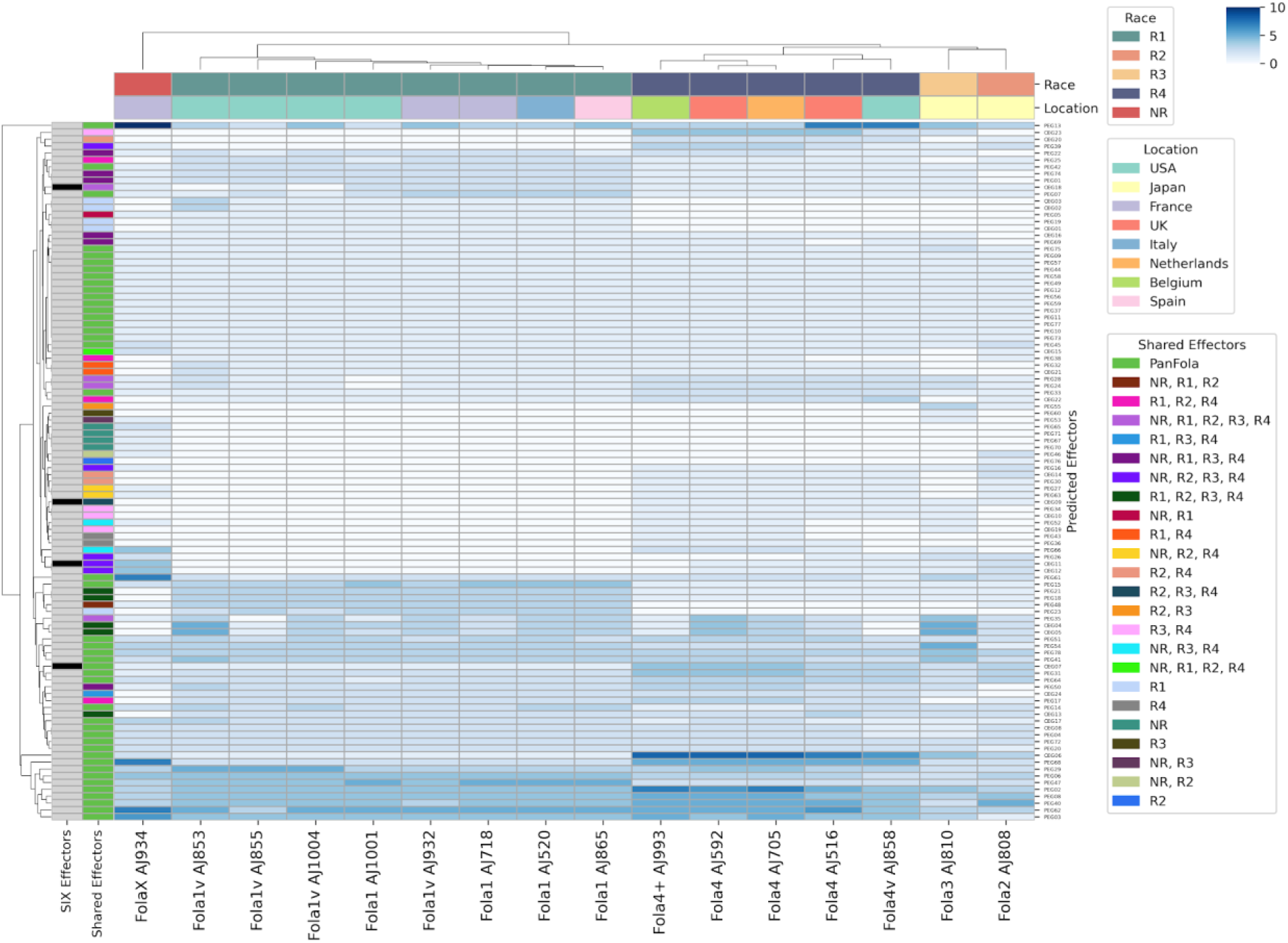
Presence/absence matrix and copy number variation of predicted effectors across Fola isolates. Heatmap showing the distribution and copy number of 102 predicted effectors (rows) across the 16 analysed Fola isolates (columns). The colour intensity represents the gene copy number (ranging from 0 to >10, as indicated by the scale). Top annotation bars denote the assigned race (Fola1 - R1, Fola2 - R2, Fola3 - R3, Fola4 - R4, NR for the novel race FolaX) and the geographic location of isolation. The left annotation bars categorise effectors based on being a SIX gene (black) and their shared distribution across the different races, e.g., PanFola (found in all 16 isolates), race-specific, or shared between specific subsets.

The number of predicted effectors varied from 72 in Fola1 AJ1001 and FolaX AJ934 to 85 in Fola4 isolates AJ592 and AJ705, with copy numbers from 1 to 17 (Fig. 3). A total of 44 (43%) were present in all genomes, including four previously identified as expressed in either Fola1 or Fola4 (CEG06, CEG07 [*SIX9*], CEG08, and CEG17); we named this core set ‘PanFola’. While 12 of these were single-copy, most exhibited copy number variations (CNVs). For instance, PEG13 was present at an average of four copies in most Fola isolates but was expanded to 17 copies in FolaX AJ934 (Fig. 3). Despite the large PanFola set, each race possessed a few unique effectors: CEG01, CEG02 and CEG03 in Fola1; PEG36 and PEG43 in Fola4; PEG76 in Fola2 (AJ808); PEG60 in Fola3 (AJ810); and four in FolaX (PEG65, PEG67, PEG70, PEG71).

The predictions for the three Fola *SIX* genes agreed with the PCR profiling (Fig. 3). CEG11 (*SIX8*) was single copy in Fola3 and Fola4, except for Fola4+ where it was absent, and present at two copies in Fola2 and four in FolaX. CEG07 (*SIX9a*) was ubiquitous but showed substantial CNVs, from one copy in Fola1 and FolaX to four copies in three Fola4 isolates (AJ993, AJ592, AJ705). CEG09 (*SIX9b*) was consistently single copy in Fola2, Fola3 and Fola4. Contrary to the PCR results, CEG18 (*SIX14*) was detected in 14 of 16 isolates, the exceptions being Fola1v AJ853 and AJ1004, where the pipeline did not identify it, suggesting a TE integration impeded automated gene detection in these two isolates.

FolaX AJ934 clustered separately from all other Fola. This isolate contained 153 predicted effectors, 72 excluding CNVs (Fig. 3), and showed a distinct profile, particularly in its CEG repertoire. Where other isolates had 15-21 CEGs, FolaX had only 9, of which three were *SIX* genes *(SIX8, SIX9a, SIX14*) and one was *PSE1*. By contrast, its 63 PEGs sat at the higher end of the observed range (56–64), 40 of them PanFola.

For the Fola1 resistance-breaking variants, no single presence/absence difference universally defined them. Instead, they showed CNVs and specific losses, clustering together through a shared expansion of PEG29 and loss of one copy of the co-localised pair PEG47 and PEG15 (Fig. 3). Relative to the standard Fola4 isolates, Fola4 AJ858 showed extensive loss and CNVs: PEG36, PEG66 and the divergently transcribed pair CEG04/CEG05 were absent; at least one copy of several PEGs (PEG02, PEG04, PEG06, PEG35, PEG40, PEG51, PEG61, PEG62) and three CEGs (CEG06, CEG08, CEG23) was lost, while an extra copy of CEG22 was gained (Fig. 3). The resistance-breaking Fola4+ variant AJ993 lost at least one copy of PEG29, PEG35, CEG04 and CEG05, plus the complete loss of CEG11 (*SIX8*) and CEG12 (*PSE1*), in agreement with Mestdagh et al. (2025). Comparison of Fola4+ AJ993 with Fola4 isolate AJ592 revealed a 70 kb deletion on an AR contig (contig_24) spanning the *SIX8/PSE1* locus and other effectors (CEG17, PEG29; Fig. 4a). Aligning Fola4+ reads against the Fola4 genome confirmed this structural variant, showing a complete loss of read coverage across the 70 kb region and a high density of split reads spanning the junction (Fig. 4b). The breakpoints were flanked by two distinct TEs, a hAT-Restless element at the 5’ end and a TcMar-Fot1 at the 3’ end, and alignment of the chimeric junction against the reference identified a 3 bp ‘GAG’ microhomology motif at the recombination site (Fig. 4c).

**Figure 4.**
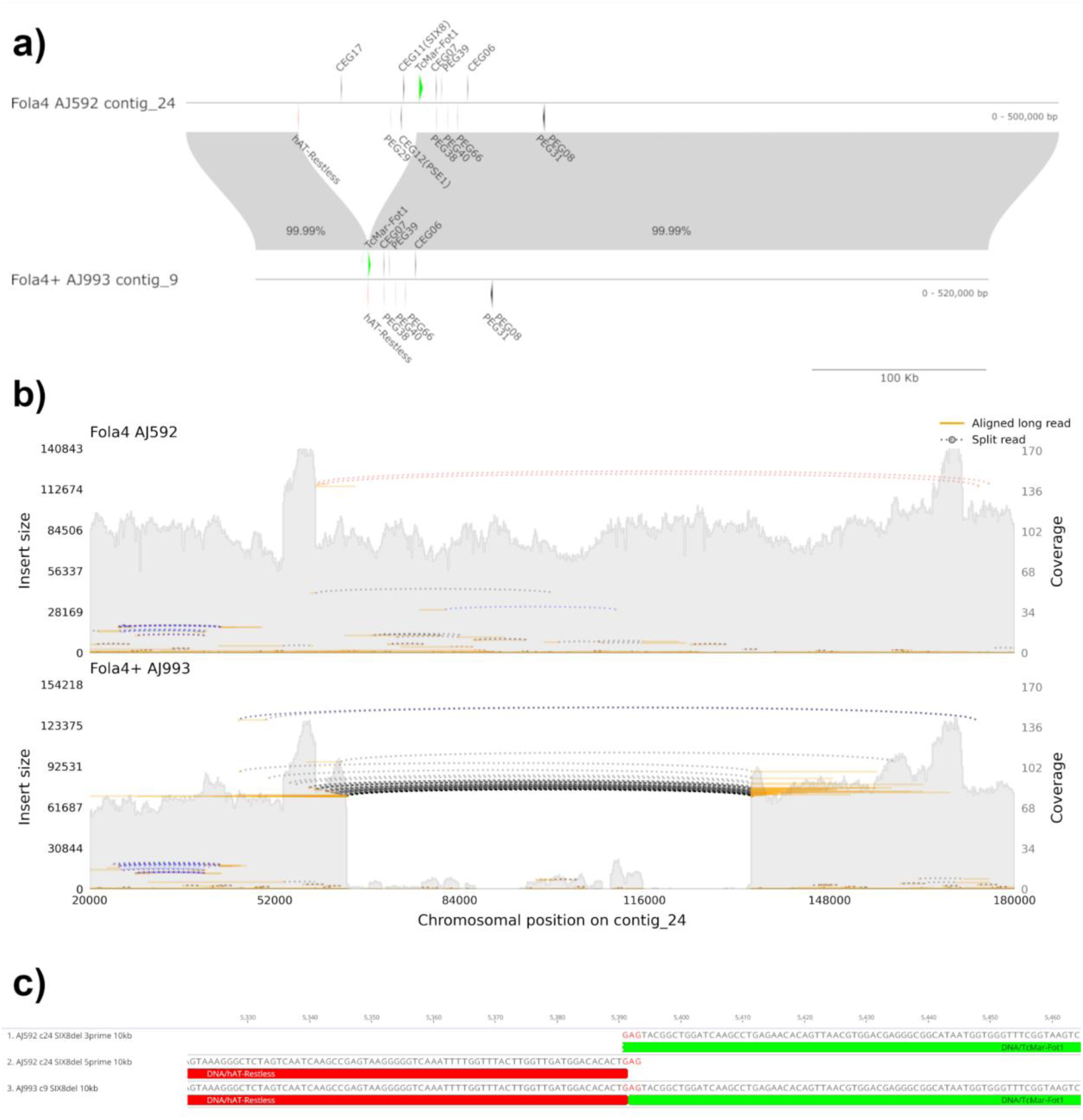
Structural variation characterising the Fola4+ (AJ993) resistance-breaking variant. (a) Synteny plot comparing AR contig 24 of Fola4 (AJ592) with contig 9 of Fola4+ (AJ993), highlighting a ∼70 kb deletion in Fola4+ that encompasses the *SIX8/PSE1* locus and other putative effectors. Transposable elements flanking the deletion region are highlighted in red (hAT-Restless) and green (TcMar-Fot1). (b) Read coverage and alignment map of Fola4 (AJ592, top) and Fola4+ (AJ993, bottom) sequencing data aligned against the Fola4 (AJ592) reference genome. The sudden drop in read coverage and the presence of split reads (dotted lines) confirm the structural deletion. (c) Local nucleotide sequence alignment at the deletion junction, revealing a 3 bp (’GAG’) microhomology motif at the site of recombination, flanked by hAT-Restless (red) and TcMar-Fot1 (green) transposable elements.

In summary, effector gene repertoires delineated the Fola races, while resistance-breaking variants were characterised by effector CNVs or specific gains/losses, most notably the complete deletion of the *SIX8/PSE1* locus in Fola4+ AJ993. FolaX was unique, with an expanded PEG complement but a notably reduced CEG repertoire, establishing it as a distinct isolate within Fola.

### Accessory region inheritance in putative new race FolaX

Despite the low inter-race AR sharing (Fig. 2b), a high proportion of effectors were found across all Fola genomes (Fig. 3). A consensus phylogeny from the 44 PanFola effector genes (Fig. 5a) was largely congruent with the core-genome phylogeny (Fig. 2b), indicating substantial vertical inheritance, though lower branch support pointed to conflicting signals. Splits networks confirmed this; the core contig network was tree-like (Fig. S8), whereas the AR network was highly reticulate, with prominent parallelograms connecting distant lineages (Fig. 5b), consistent with recombination or horizontal exchange between lineages. The apparent distinctiveness of FolaX AJ934 was mitigated by its connection into this central exchange network, suggesting its emergence may be linked to HGT with other Fola isolates.

**Figure 5.**
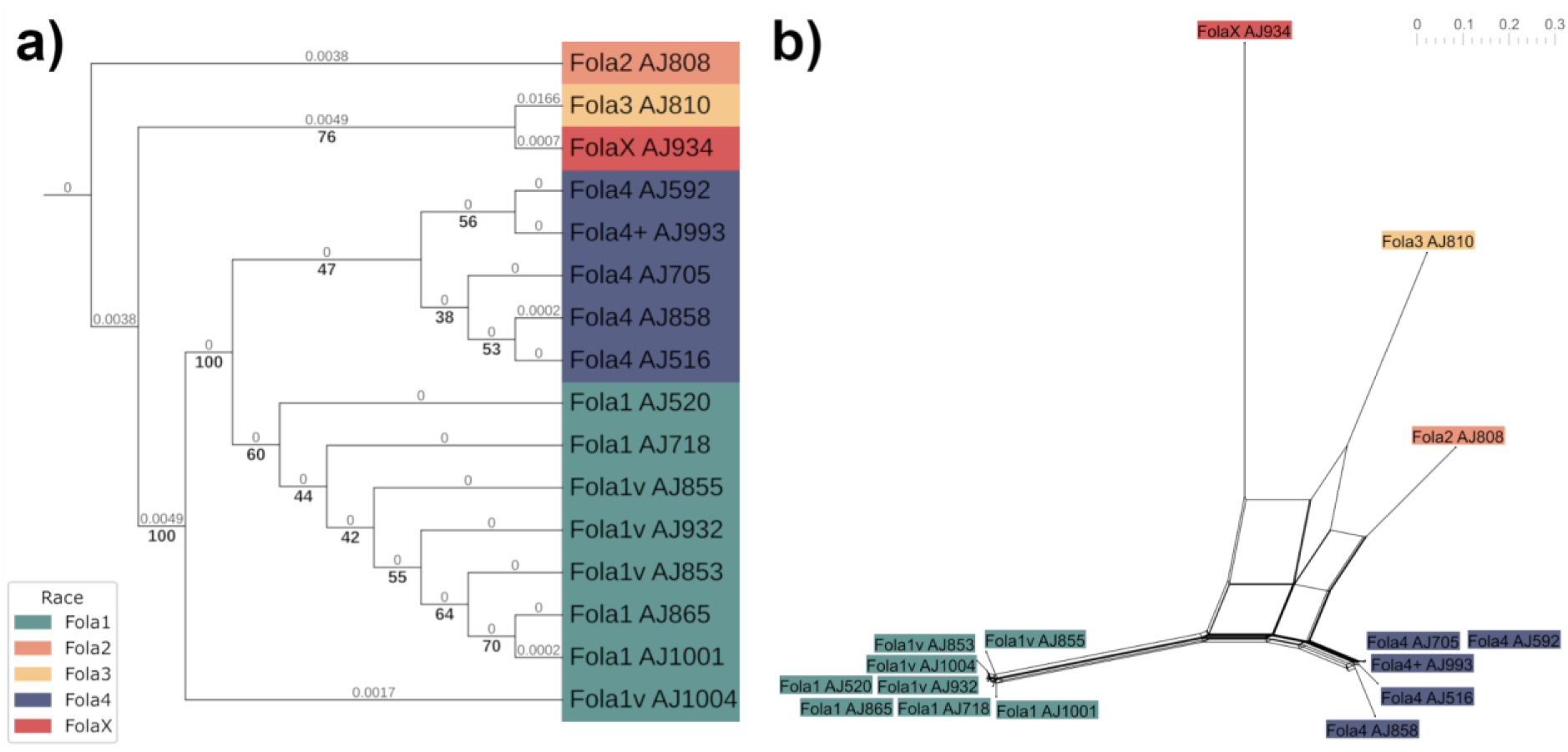
Phylogenetic inheritance and reticulate evolutionary history of Fola accessory regions (ARs). (a) Consensus maximum-likelihood phylogeny constructed from the 44 conserved PanFola effector genes. Branch lengths are indicated above the branches, and bootstrap support values are indicated below. (b) Splits network (SplitsTree) generated from the k-mer profiles of the AR contigs across all Fola isolates. The Fola races are highlighted in green (Fola1), orange (Fola2), yellow (Fola3), blue (Fola4), and red (FolaX).

Comparative k-mer analysis of FolaX identified 1,614,229 k-mers (k = 31 bp) exclusive to FOSC Clade 2 (vertical inheritance) and 894,876 shared across Fola isolates (horizontal acquisition), defining distinct trajectories across accessory contigs (Fig. S9): AR contig 24 was almost entirely shared Fola sequence, contigs 22 and 25 primarily ancestral, and the rest mosaic (Fig. 6a). Predicted effectors were significantly enriched in the shared regions (enrichment ratio = 5.56, p = 2.55 × 10⁻²⁵; 38 effectors, 32 on contigs 1 and 16) but not in vertically inherited regions (ratio = 1.007, p = 1.0). Of 94 rearrangement boundaries marking shifts between shared and ancestral sequence, regions within 5 kb were associated with TEs, predominantly TcMar-Fot1 and TcMar-Tc1. These boundary TEs had lower Kimura-80 distances than the genome average (0.182 versus 0.213), significantly so for both families (difference = −0.04; p < 0.001 and p < 0.005, respectively), suggesting more recent proliferation at recombination junctions.

**Figure 6.**
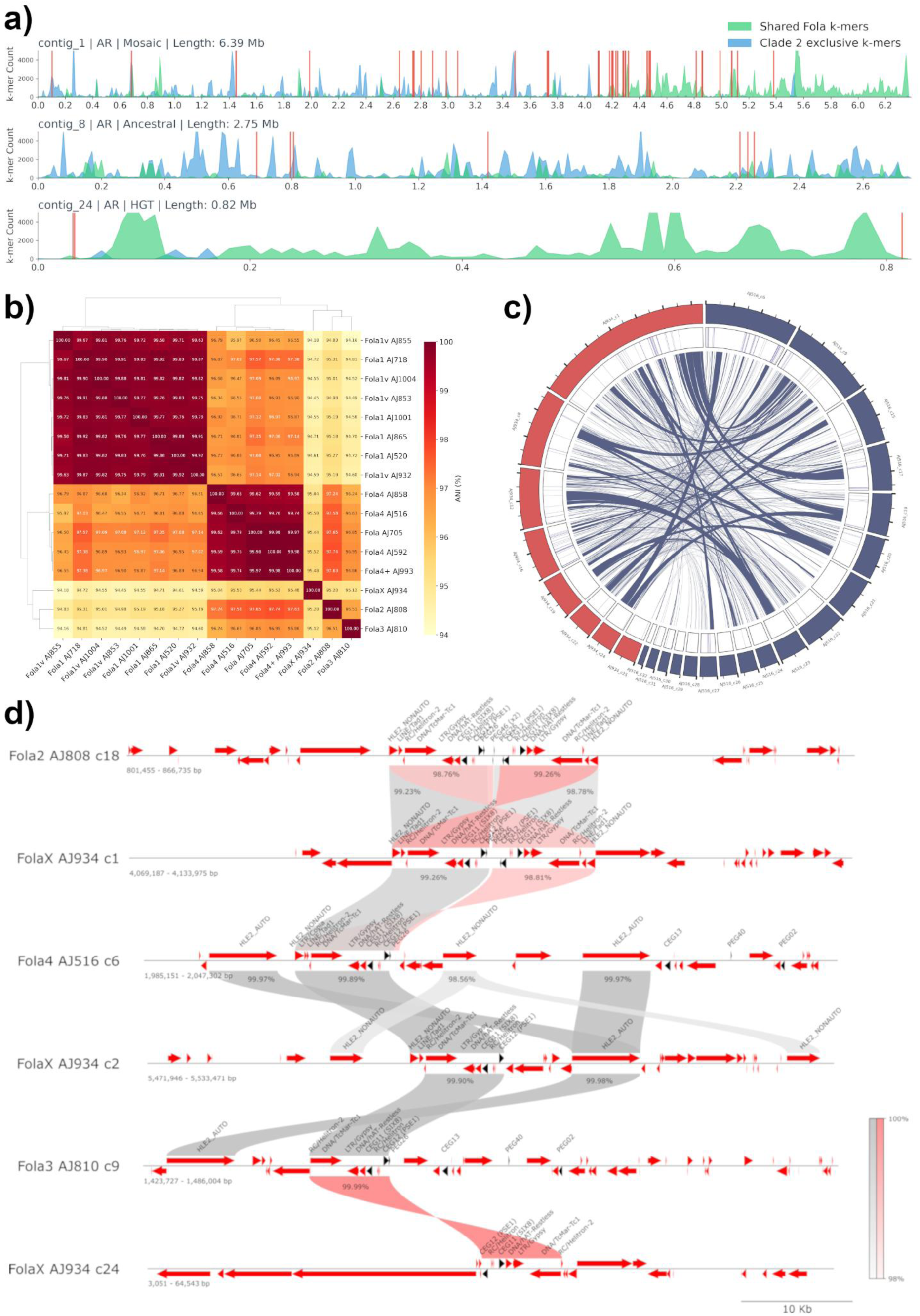
Genomic mosaicism and horizontal gene transfer (HGT) in the putative novel race FolaX (AJ934). (a) Density plots of canonical 31-mers mapped across representative FolaX AR contigs (contigs 1, 8, and 24). Green peaks indicate k-mers shared with other Fola lineages (putative HGT), whilst blue peaks indicate sequences exclusive to FOSC Clade 2 (ancestral). Red lines indicate the positions of predicted effectors. (b) Heatmap of pairwise Average Nucleotide Identity (ANI) for ARs across all of the Fola isolates. The heatmap colour scale (94 to 100) indicates the ANI percentage. (c) Circos plot detailing AR alignment between FolaX (red) and Fola4 AJ516 (blue). Syntenic regions with nucleotide identity >96% and length >5 kb are indicated by the blue ribbons. The positions of the predicted effectors are shown by the blue lines in the second track. (d) Linear comparative genomic maps of the *SIX8/PSE1* locus, showing high pairwise sequence identity (>98%) between FolaX and Fola2, Fola3, and Fola4. Black arrows represent putative effectors, and red arrows represent predicted transposable elements (TEs). Alignments in the same direction are shown in grey, whilst alignments in reverse orientation are shown in red, with colour saturation indicating the nucleotide identity.

As Starship transposons drive fungal HGT (Liu et al., 2025; Urquhart *et al*., 2025), we examined their spatial relationship with predicted effectors in FolaX. The genome contained 11 Captain genes and five Starship elements (42.6–135.9 kb); a proximity window around orphan Captain genes (tyrosine recombinases unassociated with predicted elements) accounted for fragmented or unannotated Starships. Only one effector, the FolaX-unique PEG67 (contig 4), lay within a predicted Starship (Fig. S10), and only five effectors fell within 250 kb of orphan Captain genes (four distinct: PEG14, PEG50, PEG52, PEG54; Fig. S10). K-mer density revealed a spatial disconnect between Starship machinery and the major HGT hotspots; the strongest HGT signal localised to the 3’ end of contig 1 (4.0–6.4 Mb; sixfold higher density than the 5’ region, 827.2 versus 142.6), yet this effector-dense region contained no predicted Starships or Captain genes (Fig. S10).

ANI across all Fola isolates showed FolaX core contigs were highly similar to other races (97.29% Fola2 to 98.06% Fola3; Fig. S11), whereas its AR contigs were lower (94.18% Fola1 AJ855 to 95.52% Fola4 AJ592; Fig. 6b). Whole-genome alignments of FolaX against one isolate per race (Fola1 AJ520, Fola2 AJ808, Fola3 AJ810, Fola4 AJ516), retaining syntenic regions >5 kb at >96% identity (Fig. 6c, Fig. S12), overlapped >90% of the k-mer regions, greatest with Fola3 (66.3%) and Fola4 (65.8%), then Fola2 (52%) and Fola1 (39%).

Several high-identity AR regions in FolaX contained CEGs. CEG17 was found in shared regions of 40-300 kb common to all four other-race isolates, with evidence of high-frequency transposition (Fig. S13). The divergently transcribed CEG11 (*SIX8*)/CEG12 (*PSE1*) pair lay in an 8.5-9 kb region shared with Fola2, Fola3 and Fola4 (AJ516) at >99% identity (Fig. 6d). This near-identical cassette consistently contained predicted TcMar-Tc1, hAT-Restless and Gypsy TEs, suggesting a mobility mechanism (Fig. 6d). CEG06, CEG16 and CEG18 (26–64 kb, 7.5 kb, 85 kb) were shared with Fola4 AJ516, and CEG07 (*SIX9a*)/CEG08 lay in a 13 kb region shared with Fola1 AJ520 (Table S5).

Eight of the nine FolaX CEGs (Fig. 3) lay in these high-identity regions, four in the PanFola set, and their gene trees revealed identical or near-identical sequences across the polyphyletic isolates (Fig. S14): CEG06, CEG07 (*SIX9a*), CEG17 and CEG18 (*SIX14*) clustered within the Fola clade with sequences identical to other races, while CEG08, CEG11 (*SIX8*) and CEG16 differed by two, one, and one-to-two nucleotides, respectively. A consensus tree from these shared CEGs was strongly discordant with the core genome tree (RF = 95.83%; Fig. 7). The core tree placed FolaX within FOSC Clade 2, distant from other Fola isolates, whereas the effector tree showed it deeply nested among them on very short branches. The Procrustes test confirmed low correlation (r = 0.4291, p = 0.0390), and residual analysis ranked FolaX 6th of 27 isolates (residual 0.0048), with Fola2 (AJ808; rank 1) and Fola3 (AJ810; rank 3) also high (Fig. S15).

**Figure 7.**
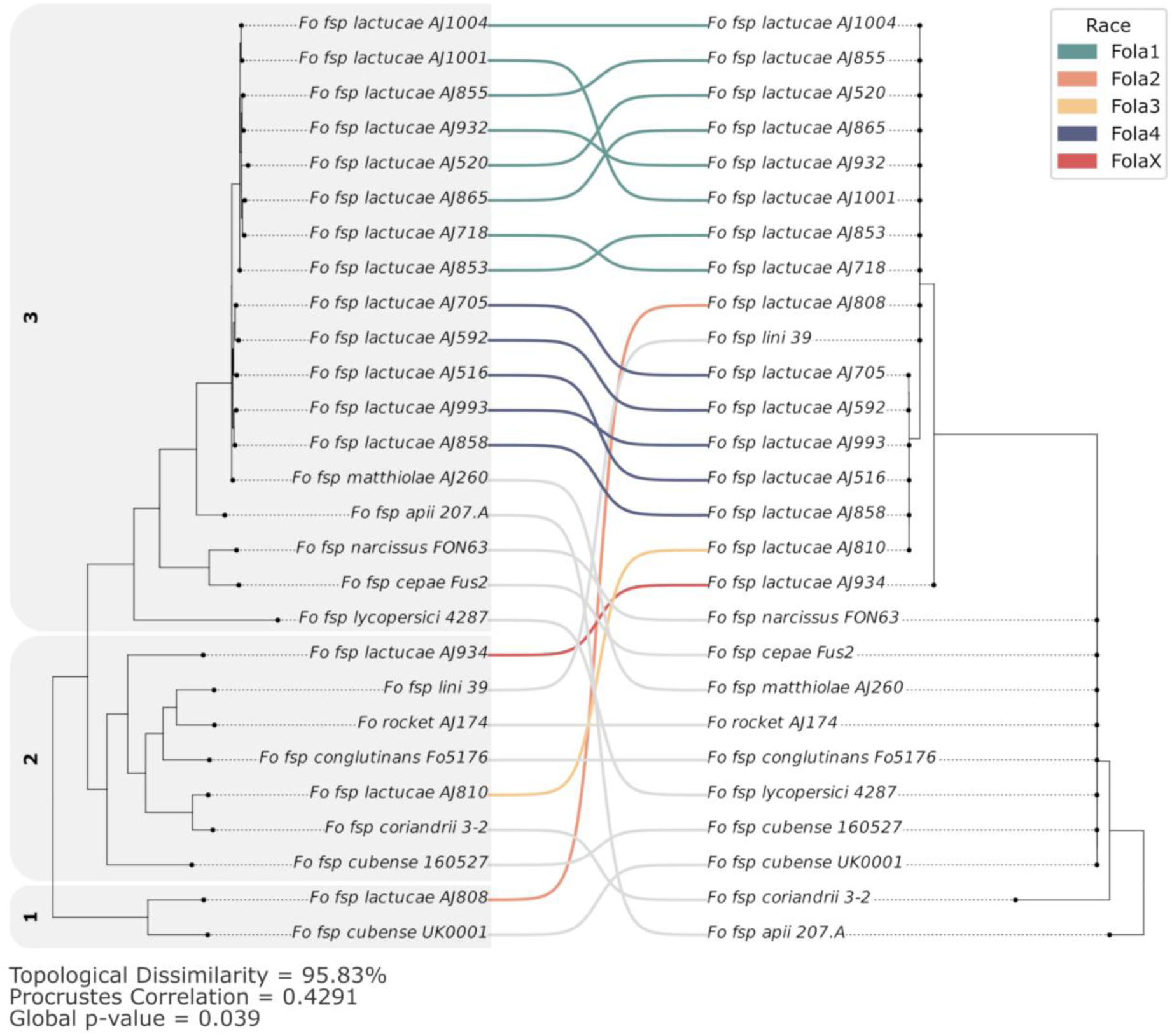
Topological incongruence between core genome and effector phylogenies. Co-phylogenetic tanglegram comparing the core genome tree based on 3,804 single-copy orthologs (left) with the consensus tree derived from eight shared candidate effector genes (right). Connecting lines map individual isolates between the two topologies. The Fola races are highlighted with coloured lines in green (Fola1), orange (Fola2), yellow (Fola3), blue (Fola4), and red (FolaX). The three FOSC clades of the core genome tree are indicated by the shaded boxes and identified by the corresponding number. The crossing lines and distinct clustering patterns visually represent a high degree of topological dissimilarity (Robinson-Foulds distance = 95.83%; Procrustes correlation = 0.429, p = 0.039), underscoring the discordance between vertical descent and horizontal effector acquisition.

Together, these findings indicate the FolaX AJ934 accessory genome is a mosaic of regions closely related to multiple Fola races. The discordance between core and effector phylogenies, the identical or near-identical effector sequences across polyphyletic isolates, and their localisation within large high-identity regions strongly support horizontal exchange of accessory genomic material in Fola.

## Discussion

The rapid emergence of resistance-breaking Fola isolates presents a significant and escalating challenge to the global lettuce industry. This study provides the first comprehensive genomic analysis of all four established races of Fola, alongside recent host resistance-breaking variants, the first genome assemblies for Fola2 and Fola3, and the characterisation of a putative new race FolaX, which overcomes multiple sources of lettuce resistance. Our analysis represents the first time whole-genome phylogenies have been used to demonstrate that Fola is polyphyletic. These results indicate that the ability to infect lettuce has emerged independently on multiple occasions, likely facilitated by the horizontal acquisition of ARs. We identified a conserved set of PanFola effectors shared across these divergent lineages, while revealing distinct evolutionary trajectories in emerging variants, ranging from CNVs in Fola1 variants to the *SIX8/PSE1* locus-specific deletion in Fola4+, and the unique molecular signature of the highly divergent FolaX. These findings extend a growing body of evidence that *F. oxysporum* ff. spp. are dynamic, reticulate networks rather than monophyletic lineages, with pathogenicity genes flowing freely across phylogenetic boundaries via mobile genetic elements (Ma *et al*., 2010; van Dam and Rep, 2017).

### Polyphyletic origins and the horizontal transfer of pathogenicity

Phylogenomic analysis of ten newly sequenced Fola isolates, combined with our previously published Fola1 and Fola4 genomes (Bates *et al*., 2024), confirms that the ability to infect lettuce reflects a reticulate evolutionary history rather than a single ancestral adaptation. While Fola1 and Fola4 share a monophyletic origin within FOSC Clade 3, Fola2, Fola3, and FolaX occupy divergent positions in Clades 1 and 2 (O’Donnell *et al*., 1998, 2009). This lack of shared core genome ancestry implies that lettuce pathogenicity has been independently acquired multiple times during FOSC evolution, most likely through horizontal transfer of pathogenicity related genes rather than convergent evolution. The polyphyletic nature of *F. oxysporum* ff. spp. has been increasingly recognised, with similar patterns reported for *F. oxysporum* f. sp. *lycopersici* (Ma *et al*., 2010), *F. oxysporum* f. sp. *cubense* (Maryani *et al*., 2019), and *F. oxysporum* f. sp. *niveum* (van Dam *et al*., 2017), suggesting host adaptation in this complex is a labile trait shaped by horizontal gene flow rather than vertical descent.

Despite this divergence, we identified a PanFola set of 44 predicted effectors present in all sequenced isolates. The near-identical nucleotide sequences of these effectors across distinct Fola clades provides strong evidence for HGT over convergent evolution. For example, *SIX8* exhibits only a single non-synonymous mutation (V88M) across Fola2, Fola3, Fola4 (AJ516), and in all four copies in FolaX, while *SIX9* and *SIX14* are identical across all races. Such sequence conservation is inconsistent with independent acquisition and strongly implies horizontal transfer from a common source or exchange network. This is also evident in other *F. oxysporum* pathosystems, where effector genes show remarkably low divergence across ff. spp. despite core genome divergence, supporting a model of recurrent horizontal transfer rather than vertical inheritance (Ma et al., 2010; van Dam et al., 2016; Henry et al., 2020, 2021).

While recent studies have linked large Starship transposons to the evolution of fungal pathogens (Peck *et al*., 2024; Sato *et al*., 2025), our analysis revealed no such association with HGT signatures in FolaX. Nevertheless, TEs have long been recognised as facilitators of genome plasticity in *F. oxysporum*, with ARs enriched for both TEs and effector genes (Ma *et al*., 2010; van Dam and Rep, 2017). The association of these effectors with TEs, including the consistent localisation of *SIX8*/*PSE1* with TcMar-Tc1, hAT-Restless, and Gypsy elements in Fola, suggests that mobile genetic elements are the primary vehicles for this exchange. The clustering of several *F. oxysporum* f. sp. *lini* effectors within Fola clades further supports a shared effector reservoir across the wider FOSC, supporting a model where host specificity is determined by the horizontally acquired effector combination, not vertical inheritance (Ma *et al*., 2010; van Dam *et al*., 2016; Henry *et al*., 2020, 2021).

### FolaX: A highly divergent putative new race

The most striking finding of this study is the identification of FolaX, a putative new race from France that is highly divergent from all established Fola lineages. Unlike the Fola1 and Fola4 variants, which represent within clade adaptations, FolaX possesses a core genome that is ancestral to a clade containing Fola3 and several non-lettuce-infecting ff. spp. This separation suggests an independent acquisition of lettuce pathogenicity, likely through complex recombination of ARs from multiple sources. With its unique molecular profile and broad virulence across the standard lettuce differential set, FolaX represents a new fundamentally different evolutionary lineage that has nonetheless assembled a functional pathogenicity toolkit. Our comparative k-mer and structural analyses support a model where the FolaX genome was assembled through segmental HCT followed by extensive, recently active TE-mediated reshuffling of ancestral and acquired components. While FolaX vertically inherited an accessory foundation from its FOSC Clade 2 common ancestor shared with Fola3, pathogenicity on lettuce appears to be largely driven by the horizontal acquisition of the Fola-specific effector pool. Despite overall AR divergence, FolaX shares regions of high nucleotide identity (>99% ANI) with other Fola races that contain key effectors, including the *SIX8*/*PSE1* locus. Notably, FolaX contains four copies of both *SIX8* and *PSE1*, compared to single copies in Fola3 and Fola4 and two copies in Fola2. This multiplication may reflect several independent HGT events or post-transfer duplication, both of which have been documented in *F. oxysporum* (van Westerhoven *et al*., 2024).

While FolaX meets the biological criteria for a new race (i.e., a unique virulence profile on lettuce differentials and a distinct genomic architecture) we refer to it as putative pending formal recognition as further work is required to determine if it arose in France or elsewhere and if it is established or widespread elsewhere in Europe. This requires multi-site surveys and validation across independent research groups and commercial breeding programmes.

The emergence of FolaX may have been driven by selection from regionally prevalent lettuce cultivars and environmental conditions as well as the presence of other Fola isolates. Fola1 and Fola4 have increasingly been found to coexist in both protected and open field environments across Europe (Gilardi *et al*., 2017b; Claerbout *et al*., 2023), and increased vigilance through the International Lettuce Fusarium Evaluation Board is identifying further putative Fola variants and races. This diverse mix provides the potential for further HGT and more rapid evolution of virulent resistance-breaking races. This is exemplified by FolaX where a mosaic of effectors from multiple Fola races were acquired through HGT, broadening its host range. Its virulence across all cultivars representing different resistance sources suggests FolaX may pose a significant threat to the lettuce industry should it become predominant in Europe.

### Race-specific mechanisms of resistance breaking

In contrast to the divergent FolaX, the lettuce resistance-breaking variants within Fola1 and Fola4 demonstrate how established races adapt through more localised genomic changes. A recurrent feature is the mobility and plasticity of the *SIX8/PSE1* locus, present in Fola2, Fola3, and Fola4, but absent in Fola1, potentially reflecting an ancestral loss or a failure in horizontal acquisition. Deletion of *SIX8* reduces Fola4 virulence in susceptible cultivars like cv. Steamboat (Legg *et al*., 2026), yet facilitates a host jump to Fola4-resistant cultivars, such as cv. Vestia, in the Fola4+ variant (AJ993; Mestdagh *et al*., 2025). This transition is characterised by a 70 kb AR deletion encompassing the entire *SIX8/PSE1* locus. Identification of a 3 bp ‘GAG’ microhomology motif at the chimeric junction suggests this rearrangement was driven by Microhomology-Mediated End Joining (MMEJ) or Non-Allelic Homologous Recombination (NAHR) between flanking transposable elements (McVey and Lee, 2008; Hoang *et al*., 2010). This parallels the loss of Avr1 (*SIX4*) observed in the emergence of Fol race 2 (Biju *et al*., 2017). Consequently, if host recognition is triggered by SIX8 or PSE1, the loss of these specific Avr genes provides a robust mechanism for evading lettuce immunity.

The genetic basis for breakdown of lettuce resistance in the US Fola1 variants AJ853, AJ932 and AJ1004 (VSP-916), appears more nuanced. These isolates were characterised by expansion of PEG29 and loss of one copy of a co-localised putative effector pair (PEG47 and PEG15), rather than large chromosomal deletions. As all CEGs retain nucleotide sequences identical to canonical Fola1, point mutations in known effectors are ruled out as the primary mechanism. Promoter changes could instead alter expression, as observed in Fola1 with the 29 bp deletion in the CEG17 promoter and the TE insertion into the *SIX14* promoter (Bates *et al*., 2024), demonstrating the need for *in planta* RNA-seq in future studies. These findings suggest that while effector loss is a powerful means of evading plant immunity, pathogens also use genomic plasticity to fine-tune effector dosage. Far less diverged from their canonical races than FolaX is from any known lineage, these variants represent stepwise, incremental adaptation rather than a major phylogenetic leap.

### Implications for breeding and diagnostics

The polyphyletic origin of Fola and the dynamic nature of its accessory genome have important implications for breeding durable resistance. The shared PanFola effector set identifies candidate targets for potential broad-spectrum resistance, and stacking R genes that recognise conserved effectors could prove more durable than targeting race-specific effectors prone to loss or modification. This has precedent in tomato, where pyramiding R genes against multiple Fol Avr genes (e.g. *I, I-2, I-3*) has given more durable resistance than single-gene approaches (Houterman *et al*., 2009; Takken and Rep, 2010). However, the emergence of FolaX and the rapid effector turnover in Fola4+ show that the Fola pangenome is highly dynamic and may circumvent resistance based solely on known effectors, making continuous surveillance of effector repertoires in emerging isolates essential.

Our data also highlight the limitations of PCR diagnostics targeting single effector genes. The *SIX8*-lacking Fola4+ variant would be misclassified as Fola1 by assays relying on *TEF1ɑ* sequence and *SIX8* presence, and structural modifications at the *SIX14* locus in Fola1 variants can give false-negative results. A multi-locus approach combining *TEF1ɑ* sequencing, race-specific markers and multiple effector profiling offers more robust identification, but still may fail to detect emerging races, as in this study for FolaX isolates, which were not amplified by race-specific primers. Ideally, whole-genome sequencing should be used for accurate race designation, particularly as new variants emerge. As Fola1 and Fola4 increasingly coexist in European production systems, the potential for further horizontal transfer and the emergence of divergent races like FolaX remains a significant threat to global lettuce production.

### Conclusions

In summary, we show that the emergence of Fola races and variants is driven by diverse genomic mechanisms, from large-scale deletions mediated by TE recombination to CNVs and the acquisition of effectors through HGT. The polyphyletic origin of Fola underscores its exceptional evolutionary plasticity, while the emergence of FolaX, characterised by a distinct effector repertoire and broad virulence spectrum highlights the continuing threat posed by this pathogen. Sustained surveillance, functional characterisation of effector genes, and breeding strategies targeting conserved virulence determinants will be critical for safeguarding global lettuce production against this rapidly evolving challenge.

## Acknowledgements

The authors gratefully acknowledge the generous contribution of Fola isolates to this project from Jim Correll and Hannah V. Zima (University of Arkansas), Kelley Richardson (USDA-ARS, Salinas), and Mathieu Pel (Enza Zaden, The Netherlands). The FolaX isolates were kindly provided by Carole Halgand (CTIFL) as part of the CASDAR project ‘Actifol’ where initial isolate phenotyping was done by project partners Bejo, Gautier, Enza Zaden, GEVES, Rijk Zwaan and Vilmorin. The authors acknowledge Research Computing at the James Hutton Institute for providing computational resources and technical support for the “UK’s Crop Diversity Bioinformatics HPC” (BBSRC grants BB/S019669/1 and BB/X019683/1), use of which has contributed to the results reported within this paper. We also acknowledge PhD funding from BBSRC (BB/X511766/1) which facilitated Fola isolate profiling and phenotyping, and funding from ‘Flanders Innovation & Entrepreneurship (VLAIO)’ (grant no. HBC.2018.2200) for Fola4+ research.

## Author contributions

RJP, EvB, HB and JC planned and designed the research. RJP conducted bioinformatics, including genome assembly and comparative analyses. EvB conducted molecular profiling and lettuce phenotyping, with assistance from SJ. HB and EV performed DNA extractions and genome sequencing. HM conducted research on Fola4+. RJP, EvB, HB, HM and JC analysed and interpreted the data. RJP, EvB, HB, MH and JC wrote and edited the manuscript. MH and JC secured funding for the project. All authors reviewed and approved the final manuscript.

## Data Availability

The datasets generated for this study can be found in the article and supplementary material. All raw sequencing data and new genome assemblies presented here have been submitted to the NCBI under the BioProject ID PRJNA1463033. Custom analysis scripts are available at https://github.com/rj-price/novel_fola_analysis with usage instructions.

## Supporting Information

**Table S1 -** Fola isolate origins, isolate codes and molecular profiling results

**Table S2 -** PCR primers used for molecular profiling

**Table S3 -** Additional *F. oxysporum* isolates used in this study

**Table S4 -** Genome assembly summary metrics and accessions

**Table S5 -** Alignment regions shared between FolaX and other Fola races with shared effectors

**Fig. S1 -** Lettuce phenotyping information and disease scales. a) Standard reactions of lettuce differential cultivars to Fola races. Exemplar Fola disease symptoms for b) wilt and c) vascular browning disease scales.

**Fig. S2 -** *F. oxysporum* TEF1ɑ amplicon phylogeny. Fola races are highlighted in pink (Fola1 & Fola4), orange (FolaX), yellow (Fola3), and purple (Fola2).

**Fig. S3 -** *F. oxysporum* SIX8 amplicon phylogeny. Fola races are highlighted in mint (SIX8a), yellow (FolaX SIX8), and green (SIX8b).

**Fig. S4 -** Wilt disease score distributions for Fola variant isolates on differential lettuce cultivars (Costa Rica No. 4, Gisela, Lomeria, Palmos and Romabella) for two experiments conducted in (a) 2024 and (b) 2025. Mean disease score distributions sharing a common letter are not significantly different by Kruskal-Wallis with Dunn’s post hoc test with Benjamini-Hochberg correction at the 5% significance level.

**Fig. S5 -** Average number of alignments to each contig of 29 *F. oxysporum* genome assemblies using a pangenomic approach based on all-vs-all whole-genome alignments. Contigs were classified as accessory where they aligned to fewer than 15 of the other *F. oxysporum* genomes (red dotted line). Host species are indicated by colour.

**Fig. S6 –** Principal component analysis of contig features across all Fola assemblies, using average number of alignments to other genomes (AverageCount), GC percentage (GC Percentage), repeat content (RepeatPercentage) and sequence data coverage per contig (Coverage). Core contigs (green), accessory region (AR) contigs (pink), and mitochondrial contigs (blue) are separated by their feature profiles. Contig length is represented by the size of the marker.

**Fig. S7 -** Total predicted effector density on core (green) and accessory (purple) contigs across Fola assemblies.

**Fig. S8 -** Splits network (SplitsTree) generated from the k-mer profiles of core contigs across all Fola isolates.

**Fig. S9 -** Comparative k-mer profiling across the FolaX genome. The Fola-specific (green) and lineage-specific (blue) k-mer sequences were mapped to the FolaX genome. Predicted effectors represented by vertical red lines.

**Fig. S10 -** Distribution of Starships and effectors in FolaX. Starship elements are represented by orange rectangles, Captain genes by red diamonds, and predicted effectors by vertical red lines. Fola-specific (green) and lineage-specific (blue) k-mer profiles are above each contig.

**Fig. S11 -** Heatmap of pairwise average nucleotide identities (ANI) of all core contigs across Fola genomes.

**Fig. S12 -** Alignment of FolaX ARs (red) to other Fola races Circular alignment plots comparing FolaX AR contigs against the AR contigs of the other four races, Fola1 (AJ520, green), Fola2 (AJ808, orange), Fola3 (AJ810, yellow), and Fola4 (AJ516, dark blue). The ribbons represent regions of high (>96%) sequence similarity, longer than 5 kb.

**Fig. S13 -** Linear comparative genomic map showing conserved regions containing CEG17 between FolaX and the other Fola races. Black arrows represent putative effectors and red arrows represent predicted transposable elements (TEs). Grey ribbons represent alignment blocks in the same orientation, red ribbons represent alignments in the opposite orientation.

**Fig. S14 -** Phylogenetic trees for candidate effector genes (CEGs) found in shared alignment regions in FolaX.

**Fig. S15 -** Phylogenetic incongruence residuals (PACo) for *F. oxysporum* isolates. The chart measures the squared residual between core and effector placement.

## References

van Amsterdam, S., Jenkins, S. and Clarkson, J.P. (2023) ‘First Report of Fusarium oxysporum f. sp. lactucae Race 1 Causing Lettuce Wilt in Northern Ireland’, Plant Disease, 0(0), p. PDIS-01-23-0196-PDN. Available at: 10.1094/pdis-01-23-0196-pdn.

Baril, T., Galbraith, J. and Hayward, A. (2024) ‘Earl Grey: A Fully Automated User-Friendly Transposable Element Annotation and Analysis Pipeline’, Molecular Biology and Evolution. Edited by I. Arkhipova, 41(4), p. msae068. Available at: 10.1093/molbev/msae068.

Bates, H.J. et al. (2024) ‘Comparative genomics and transcriptomics reveal differences in effector complement and expression between races of Fusarium oxysporum f.sp. lactucae’, Frontiers in Plant Science, 15, p. 1415534. Available at: 10.3389/fpls.2024.1415534.

Belyeu, J.R. et al. (2021) ‘Samplot: a platform for structural variant visual validation and automated filtering’, Genome Biology, 22(1), p. 161. Available at: 10.1186/s13059-021-02380-5.

Biju, V.C., et al. (2017) ‘Multiple Evolutionary Trajectories Have Led to the Emergence of Races in Fusarium oxysporum f. sp. lycopersici’, Applied and Environmental Microbiology. Edited by D. Cullen, 83(4), pp. e02548–16. Available at: 10.1128/AEM.02548-16.

Brenes Guallar, M.A., et al. (2022) ‘Fusarium oxysporum effector clustering version 2: An updated pipeline to infer host range’, Frontiers in Plant Science, 13. Available at: 10.3389/fpls.2022.1012688.

Bürkner, P.-C. et al. (2025) ‘brms: Bayesian Regression Models using “Stan”’. Available at: https://cran.r-project.org/web/packages/brms/index.html (Accessed: 2 July 2026).

Cabral, C.S. et al. (2019) ‘A single dominant gene/locus model for control of Fusarium oxysporum f. sp. lactucae race 1 resistance in lettuce (Lactuca sativa)’, Euphytica, 215(7), p. 114. Available at: 10.1007/s10681-019-2441-2.

Camacho, C. et al. (2009) ‘BLAST+: architecture and applications’, BMC Bioinformatics, 10(1), p. 421. Available at: 10.1186/1471-2105-10-421.

Chen, Y. et al. (2021) ‘Efficient assembly of nanopore reads via highly accurate and intact error correction’, Nature Communications, 12, p. 60. Available at: 10.1038/s41467-020-20236-7.

Claerbout, J. et al. (2018) ‘First Report of *Fusarium oxysporum* f. sp. *lactucae* Race 4 on Lettuce in Belgium’, Plant Disease, 102(5), p. 1037. Available at: 10.1094/PDIS-10-17-1627-PDN.

Claerbout, J. et al. (2023) ‘Fusarium isolates from Belgium causing wilt in lettuce show genetic and pathogenic diversity’, Plant Pathology, 72(3), pp. 593–609. Available at: 10.1111/ppa.13668.

van Dam, P. et al. (2016) ‘Effector profiles distinguish formae speciales of Fusarium oxysporum’, Environmental Microbiology, 18(11), pp. 4087–4102.

van Dam, P. et al. (2017) ‘A mobile pathogenicity chromosome in Fusarium oxysporum for infection of multiple cucurbit species’, Scientific Reports, 7(1), p. 9042. Available at: 10.1038/s41598-017-07995-y.

van Dam, P. and Rep, M. (2017) ‘The Distribution of Miniature Impala Elements and SIX Genes in the Fusarium Genus is Suggestive of Horizontal Gene Transfer’, Journal of Molecular Evolution, 85(1–2), pp. 14–25. Available at: 10.1007/s00239-017-9801-0.

Danecek, P. et al. (2021) ‘Twelve years of SAMtools and BCFtools’, GigaScience, 10(2), p. giab008. Available at: 10.1093/gigascience/giab008.

De Coster, W. et al. (2018) ‘NanoPack: visualizing and processing long-read sequencing data’, Bioinformatics, 34(15), pp. 2666–2669. Available at: 10.1093/bioinformatics/bty149.

Dixon, Philip (2003) ‘VEGAN, a package of R functions for community ecology’, Journal of Vegetation Science, 14(6), pp. 927–930. Available at: 10.1111/j.1654-1103.2003.tb02228.x.

Formenti, G., et al. (2022) ‘Gfastats: conversion, evaluation and manipulation of genome sequences using assembly graphs’, Bioinformatics. Edited by A. Valencia, 38(17), pp. 4214–4216. Available at: 10.1093/bioinformatics/btac460.

Fujinaga, M. et al. (2014) ‘PCR-based Race Differentiation of Fusarium oxysporum f. sp. lactucae’. Available at: https://www.semanticscholar.org/paper/PCR-based-Race-Differentiation-of-Fusarium-f.-sp.-Fujinaga-Yamagishi/db976c56030d278f7a6065bc849f62421b640ac7 (Accessed: 2 July 2026).

Gálvez, L. et al. (2023) ‘First Report of Fusarium oxysporum f. sp. lactucae Race 4 Causing Lettuce Wilt in Spain’, Plant Disease, 107(8), p. 2549. Available at: 10.1094/PDIS-12-22-2819-PDN.

Garibaldi, A., Gilardi, G. and Gullino, M.L. (2004) ‘Seed transmission of Fusarium oxysporum f.sp.lactucae’, Phytoparasitica, 32(1), pp. 61–65. Available at: 10.1007/BF02980861.

Gawehns, F. et al. (2014) ‘The Fusarium oxysporum Effector Six6 Contributes to Virulence and Suppresses I-2-Mediated Cell Death’, Molecular Plant-Microbe Interactions, 27(4), pp. 336–348. Available at: 10.1094/MPMI-11-13-0330-R.

Gilardi, G., Ortega, S.F., et al. (2017a) ‘A new race of Fusarium oxysporum f. sp. lactucae of lettuce’, Plant Pathology, 66(4), pp. 677–688. Available at: 10.1111/ppa.12616.

Gilardi, G., Pons, C., et al. (2017b) ‘Presence of Fusarium Wilt, Incited by Fusarium oxysporum f.sp. lactucae, on Lettuce in France’, Plant Disease, 101(6), p. 1053. Available at: 10.1094/PDIS-12-16-1815-PDN.

Gilardi, G. et al. (2019) ‘First Report of Fusarium oxysporum f. sp. lactucae Race 4 on Lettuce in Italy’, Plant Disease, 103(10), pp. 2680–2680. Available at: 10.1094/pdis-05-19-0902-pdn.

Gluck-Thaler, E. and Vogan, A.A. (2024) ‘Systematic identification of cargo-mobilizing genetic elements reveals new dimensions of eukaryotic diversity’, Nucleic Acids Research, 52(10), pp. 5496–5513. Available at: 10.1093/nar/gkae327.

Guerrero, M.M. et al. (2020) ‘First Report of Fusarium Wilt of Lettuce Caused by Fusarium oxysporum f. sp. lactucae Race 1 in Spain’, Plant Disease, 104(6), pp. 1858–1858. Available at: 10.1094/pdis-10-19-2143-pdn.

Halgand, C. et al. (2024) ACTIFOL Project: Knowledge acquired between 2020 and 2023. 401. INFOCTIFL. Available at: https://www.ctifl.fr/projet-actifol-connaissances-acquises-entre-2020-et-2023-infos-ctifl-401 (Accessed: 2 July 2025).

Henry, P. et al. (2020) ‘Genomic differences between the new Fusarium oxysporum f. sp. apii (Foa) race 4 on celery, the less virulent Foa races 2 and 3, and the avirulent on celery f. sp. coriandrii’, BMC Genomics, 21(1), p. 730. Available at: 10.1186/s12864-020-07141-5.

Henry, P.M. et al. (2021) ‘Horizontal chromosome transfer and independent evolution drive diversification in *Fusarium oxysporum* f. sp. *fragariae*’, New Phytologist, 230(1), pp. 327–340. Available at: 10.1111/nph.17141.

Herrero, M.L., Nagy, N.E. and Solheim, H. (2021) ‘First Report of Fusarium oxysporum f. sp. lactucae Race 1 Causing Fusarium Wilt of Lettuce in Norway’, Plant Disease, 105(8), p. 2239. Available at: 10.1094/pdis-01-21-0134-pdn.

Hoang, M.L. et al. (2010) ‘Competitive Repair by Naturally Dispersed Repetitive DNA during Non-Allelic Homologous Recombination’, PLoS Genetics, 6(12), p. e1001228. Available at: 10.1371/journal.pgen.1001228.

Houterman, P.M. et al. (2007) ‘The mixed xylem sap proteome of Fusarium oxysporum-infected tomato plants’, Molecular Plant Pathology, 8(2), pp. 215–221. Available at: 10.1111/j.1364-3703.2007.00384.x.

Houterman, P.M. et al. (2009) ‘The effector protein Avr2 of the xylem-colonizing fungus Fusarium oxysporum activates the tomato resistance protein I-2 intracellularly’, The Plant Journal, 58(6), pp. 970–978. Available at: 10.1111/j.1365-313X.2009.03838.x.

Houterman, P.M., Cornelissen, B.J.C. and Rep, M. (2008) ‘Suppression of Plant Resistance Gene-Based Immunity by a Fungal Effector’, PLoS Pathogens, 4(5), p. e1000061. Available at: 10.1371/journal.ppat.1000061.

Huson, D.H. and Bryant, D. (2024) ‘The SplitsTree App: interactive analysis and visualization using phylogenetic trees and networks’, Nature Methods, 21(10), pp. 1773–1774. Available at: 10.1038/s41592-024-02406-3.

Hutchinson, M.C., et al. (2017) ‘paco: implementing Procrustean Approach to Cophylogeny in R’, Methods in Ecology and Evolution. Edited by R. Fitzjohn, 8(8), pp. 932–940. Available at: 10.1111/2041-210X.12736.

Inami, K. et al. (2012) ‘A Genetic Mechanism for Emergence of Races in Fusarium oxysporum f. sp. lycopersici: Inactivation of Avirulence Gene AVR1 by Transposon Insertion’, PLoS ONE, 7(8), p. e44101. Available at: 10.1371/journal.pone.0044101.

International Seed Federation (2022) Final report EG ISF DRT and IBEB project Lettuce Fusarium oxysporum f. sp. lactucae (Fol: 4) resistance. Available at: https://worldseed.org/document/isf-final-report-fol-race-4-lettuce/ (Accessed: 16 November 2025).

International Seed Federation (2024) Differential Sets Fusarium oxysporum f. sp. lactucae (Fol) – Lettuce. Available at: https://worldseed.org/wp-content/uploads/2024/02/Feb-2024-Differential-sets-Fol-race4_final.pdf (Accessed: 17 June 2025).

Jain, C. et al. (2018) ‘High throughput ANI analysis of 90K prokaryotic genomes reveals clear species boundaries’, Nature Communications, 9(1), p. 5114. Available at: 10.1038/s41467-018-07641-9.

Jangir, P. et al. (2021) ‘Secreted in Xylem Genes: Drivers of Host Adaptation in Fusarium oxysporum’, Frontiers in Plant Science, 12, p. 628611. Available at: 10.3389/fpls.2021.628611.

Kalyaanamoorthy, S. et al. (2017) ‘ModelFinder: fast model selection for accurate phylogenetic estimates’, Nature Methods, 14(6), pp. 587–589. Available at: 10.1038/nmeth.4285.

Kolmogorov, M. et al. (2019) ‘Assembly of long, error-prone reads using repeat graphs’, Nature Biotechnology, 37(5), pp. 540–546. Available at: 10.1038/s41587-019-0072-8.

Kurtz, S. et al. (2004) ‘Versatile and open software for comparing large genomes’, Genome Biology, 5(2), p. R12. Available at: 10.1186/gb-2004-5-2-r12.

Langmead, B. and Salzberg, S.L. (2012) ‘Fast gapped-read alignment with Bowtie 2’, Nature methods, 9(4), pp. 357–359. Available at: 10.1038/nmeth.1923.

Legg, A.D. et al. (2026) ‘The Effector *SIX8* Contributes to Virulence of *Fusarium oxysporum* f. sp. *lactucae* Race 4 on Lettuce’, Molecular Plant Pathology, 27(6), p. e70296. Available at: 10.1111/mpp.70296.

Letunic, I. and Bork, P. (2021) ‘Interactive Tree Of Life (iTOL) v5: an online tool for phylogenetic tree display and annotation’, Nucleic acids research, 49(W1), pp. W293–W296.

Li, H. (2018) ‘Minimap2: pairwise alignment for nucleotide sequences’, Bioinformatics, 34(18), pp. 3094–3100. Available at: 10.1093/bioinformatics/bty191.

Liu, F., Cheewangkoon, R. and Zhao, R.-L. (2025) ‘Discovery of a New Starship Transposon Driving the Horizontal Transfer of the ToxA Virulence Gene in Alternaria ventricosa’, Microorganisms, 13(2), p. 376. Available at: 10.3390/microorganisms13020376.

Ma, L. et al. (2015) ‘The AVR2–SIX5 gene pair is required to activate I-2-mediated immunity in tomato’, New Phytologist, 208(2), pp. 507–518. Available at: 10.1111/nph.13455.

Ma, L.-J. et al. (2010) ‘Comparative genomics reveals mobile pathogenicity chromosomes in Fusarium’, Nature, 464(7287), pp. 367–373.

Marçais, G. and Kingsford, C. (2011) ‘A fast, lock-free approach for efficient parallel counting of occurrences of k-mers’, Bioinformatics, 27(6), pp. 764–770. Available at: 10.1093/bioinformatics/btr011.

Maryani, N. et al. (2019) ‘Phylogeny and genetic diversity of the banana Fusarium wilt pathogen *Fusarium oxysporum* f. sp. *cubense* in the Indonesian centre of origin’, Studies in Mycology, 92, pp. 155–194. Available at: 10.1016/j.simyco.2018.06.003.

McGowan, J. (2025) ‘BUSCO_phylogenomics’. Available at: https://github.com/jamiemcg/BUSCO_phylogenomics.

McVey, M. and Lee, S.E. (2008) ‘MMEJ repair of double-strand breaks (director’s cut): deleted sequences and alternative endings’, Trends in genetics : TIG, 24(11), pp. 529–538. Available at: 10.1016/j.tig.2008.08.007.

Mestdagh, H. et al. (2023) ‘Detection of Fusarium oxysporum f.sp. lactucae race 1 and 4 via race-specific real-time PCR and target enrichment’, Frontiers in Plant Science, 14. Available at: 10.3389/fpls.2023.1272136.

Mestdagh, H. et al. (2025) ‘Cultivar Resistance and Alternative Crops to Manage Fusarium Wilt in Lettuce Caused by *Fusarium oxysporum* f. sp. *Lactucae* Race 4 and the Emerging Race 4+’, Journal of Phytopathology, 173(3), p. e70072. Available at: 10.1111/jph.70072.

Mori, H. (2022) ‘pyCircos’. Available at: https://github.com/ponnhide/pyCircos.

Motohashi, S. (1960) ‘Occurrence of lettuce root rot’, Annals of the Phytopathological Society of Japan, 25, p. 47.

Murray, J.J. et al. (2021) ‘Genetic Resistance of Lactuca spp. against Fusarium oxysporum f. sp. lactucae Race 1’, HortScience, 56(12), pp. 1552–1564. Available at: 10.21273/HORTSCI16186-21.

Nayak, S. et al. (2025) ‘Detection of novel pathogenic variants of *Fusarium oxysporum* f. sp. *lactucae* in California’, Plant Pathology, 74(2), pp. 295–307. Available at: 10.1111/ppa.14019.

Nguyen, L.-T. et al. (2015) ‘IQ-TREE: A Fast and Effective Stochastic Algorithm for Estimating Maximum-Likelihood Phylogenies’, Molecular Biology and Evolution, 32(1), pp. 268–274. Available at: 10.1093/molbev/msu300.

O’Donnell, K. et al. (1998) ‘Multiple evolutionary origins of the fungus causing Panama disease of banana: Concordant evidence from nuclear and mitochondrial gene genealogies’, Proceedings of the National Academy of Sciences, 95(5), pp. 2044–2049. Available at: 10.1073/pnas.95.5.2044.

O’Donnell, K. et al. (2009) ‘A two-locus DNA sequence database for typing plant and human pathogens within the Fusarium oxysporum species complex’, Fungal Genetics and Biology, 46(12), pp. 936–948. Available at: 10.1016/j.fgb.2009.08.006.

Oxford Nanopore Technologies (2023) ‘Medaka’. Available at: https://github.com/nanoporetech/medaka.

Paradis, E., Claude, J. and Strimmer, K. (2004) ‘APE: Analyses of Phylogenetics and Evolution in R language’, Bioinformatics, 20(2), pp. 289–290. Available at: 10.1093/bioinformatics/btg412.

Pasquali, M. et al. (2007) ‘Identification of Race 1 of Fusarium oxysporum f. sp. lactucae on Lettuce by Inter-Retrotransposon Sequence-Characterized Amplified Region Technique’, Phytopathology®, 97(8), pp. 987–996. Available at: 10.1094/PHYTO-97-8-0987.

Peck, L.D. et al. (2024) ‘Horizontal transfers between fungal Fusarium species contributed to successive outbreaks of coffee wilt disease’, PLOS Biology, 22(12), p. e3002480. Available at: 10.1371/journal.pbio.3002480.

Quinlan, A.R. and Hall, I.M. (2010) ‘BEDTools: a flexible suite of utilities for comparing genomic features’, Bioinformatics, 26(6), pp. 841–842.

R Core Team (2025) ‘R: A Language and Environment for Statistical Computing.’ R Foundation for Statistical Computing, Vienna, Austria. Available at: https://www.R-project.org/.

Ranallo-Benavidez, T.R., Jaron, K.S. and Schatz, M.C. (2020) ‘GenomeScope 2.0 and Smudgeplot for reference-free profiling of polyploid genomes’, Nature Communications, 11(1), p. 1432. Available at: 10.1038/s41467-020-14998-3.

Rempel, A. and Wittler, R. (2021) ‘SANS serif: alignment-free, whole-genome-based phylogenetic reconstruction’, Bioinformatics. Edited by R. Schwartz, 37(24), pp. 4868–4870. Available at: 10.1093/bioinformatics/btab444.

Ren, J. and Chaisson, M.J.P. (2021) ‘lra: A long read aligner for sequences and contigs’, PLOS Computational Biology, 17(6), p. e1009078. Available at: 10.1371/journal.pcbi.1009078.

Rep, M. et al. (2004) ‘A small, cysteine-rich protein secreted by Fusarium oxysporum during colonization of xylem vessels is required for I-3-mediated resistance in tomato’, Molecular Microbiology, 53(5), pp. 1373–1383. Available at: 10.1111/j.1365-2958.2004.04177.x.

Revell, L.J. (2012) ‘phytools: an R package for phylogenetic comparative biology (and other things)’, Methods in Ecology and Evolution, 3(2), pp. 217–223. Available at: 10.1111/j.2041-210X.2011.00169.x.

Rhie, A. et al. (2020) ‘Merqury: reference-free quality, completeness, and phasing assessment for genome assemblies’, Genome Biology, 21(1), p. 245. Available at: 10.1186/s13059-020-02134-9.

Sato, Y. et al. (2025) ‘Starship giant transposons dominate plastic genomic regions in a fungal plant pathogen and drive virulence evolution’, Nature Communications, 16, p. 6806. Available at: 10.1038/s41467-025-61986-6.

Scott, J.C., Kirkpatrick, S.C. and Gordon, T.R. (2010) ‘Variation in susceptibility of lettuce cultivars to fusarium wilt caused by Fusarium oxysporum f.sp. lactucae’, Plant Pathology, 59(1), pp. 139–146. Available at: 10.1111/j.1365-3059.2009.02179.x.

Seki, K. et al. (2022) ‘Discovery and fine mapping of a novel resistance locus to Fusarium wilt race 2 in lettuce (Lactuca sativa L.)’, Euphytica, 218(8), p. 120. Available at: 10.1007/s10681-022-03058-0.

Shimoyama, Y. (2024) ‘pyGenomeViz: A genome visualization python package for comparative genomics’. Available at: https://github.com/moshi4/pyGenomeViz.

Simão, F.A. et al. (2015) ‘BUSCO: assessing genome assembly and annotation completeness with single-copy orthologs’, Bioinformatics, 31(19), pp. 3210–3212. Available at: 10.1093/bioinformatics/btv351.

Takken, F. and Rep, M. (2010) ‘The arms race between tomato and Fusarium oxysporum’, Molecular Plant Pathology, 11(2), pp. 309–314. Available at: 10.1111/j.1364-3703.2009.00605.x.

Taylor, A. et al. (2016) ‘Identification of pathogenicity-related genes in Fusarium oxysporum f. sp. cepae’, Molecular Plant Pathology, 17(7), pp. 1032–1047.

Taylor, A., Jackson, A.C. and Clarkson, J.P. (2019) ‘First Report of Fusarium oxysporum f. sp. lactucae Race 4 Causing Lettuce Wilt in England and Ireland’, Plant Disease, 103(5), pp. 1033–1033. Available at: 10.1094/PDIS-10-18-1858-PDN.

Urquhart, A.S., Forsythe, A. and Vogan, A.A. (2025) ‘Are Fungal Disease Outbreaks Instigated by *Starship* Transposons?’, Molecular Plant Pathology, 26(7), p. e70124. Available at: 10.1111/mpp.70124.

Vaser, R. et al. (2017) ‘Fast and accurate de novo genome assembly from long uncorrected reads’, Genome Research, 27(5), pp. 737–746. Available at: 10.1101/gr.214270.116.

Waqas, M. et al. (2025) ‘Host Range and Molecular Characterization of the Four Races of Fusarium oxysporum f. sp. lactucae’, Plant Disease, 108(7). Available at: https://apsjournals.apsnet.org/doi/10.1094/PDIS-07-24-1505-RE.

van Westerhoven, A.C. et al. (2024) ‘Segmental duplications drive the evolution of accessory regions in a major crop pathogen’, New Phytologist, 242(2), pp. 610–625. Available at: 10.1111/nph.19604.

Wick, Ryan (2018) ‘Porechop’. Available at: https://github.com/rrwick/Porechop.

Wick, Ryan (2021) ‘Filtlong’. Available at: https://github.com/rrwick/Filtlong.

Wood, D.E., Lu, J. and Langmead, B. (2019) ‘Improved metagenomic analysis with Kraken 2’, Genome Biology, 20(1), p. 257. Available at: 10.1186/s13059-019-1891-0.

Yu, H. et al. (2023) ‘Accessory Chromosomes of the Fusarium oxysporum Species Complex and Their Contribution to Host Niche Adaptation’, in B. Scott and C. Mesarich (eds) Plant Relationships: Fungal-Plant Interactions. Cham: Springer International Publishing, pp. 371–388. Available at: 10.1007/978-3-031-16503-0_16.

Zhang, C., Nielsen, R. and Mirarab, S. (2025) ‘ASTER: A Package for Large-Scale Phylogenomic Reconstructions’, Molecular Biology and Evolution, 42(8), p. msaf172. Available at: 10.1093/molbev/msaf172.

