## Supplementary material for "Polyphyletic origin and horizontal exchange drive the emergence of a novel resistance-breaking *Fusarium oxysporum* f. sp. *lactucae* race": Figure S1

**a)**

| Lettuce cv. | Fola1 | Fola2 | Fola3 | Fola4 |
|---|---|---|---|---|
| Gisela | S | S | S | S |
| Costa Rica N°4 | HR | S | S | S |
| Lomeria | S | HR | HR | HR |
| Romabella | HR | HR | S | IR |
| Palmos | HR | S | IR | HR |

**b)**

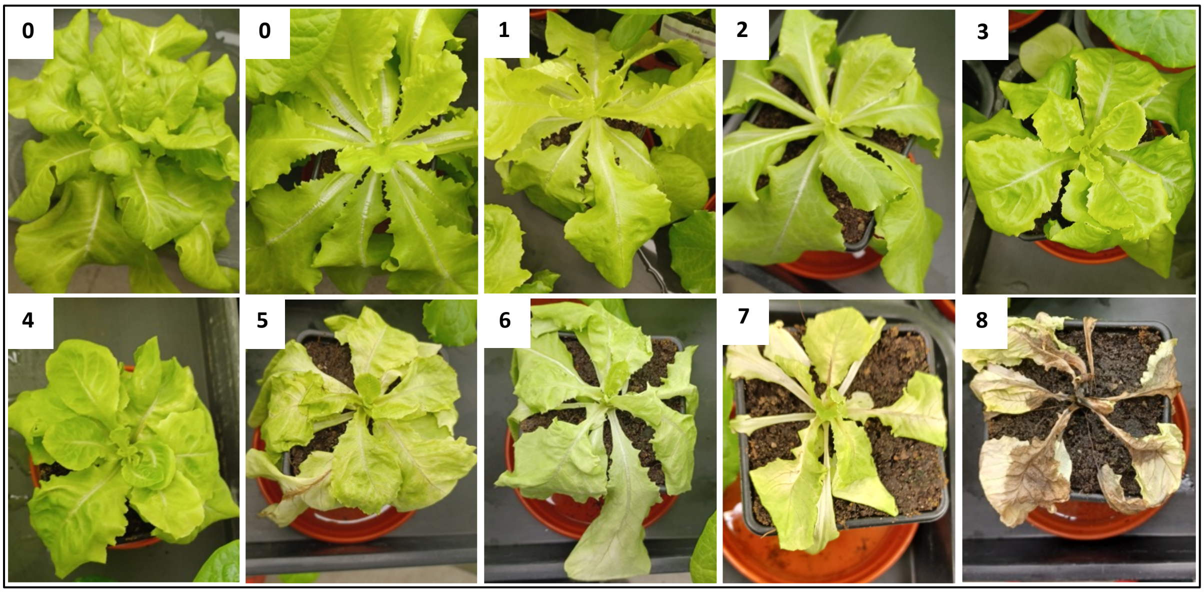


**c)**

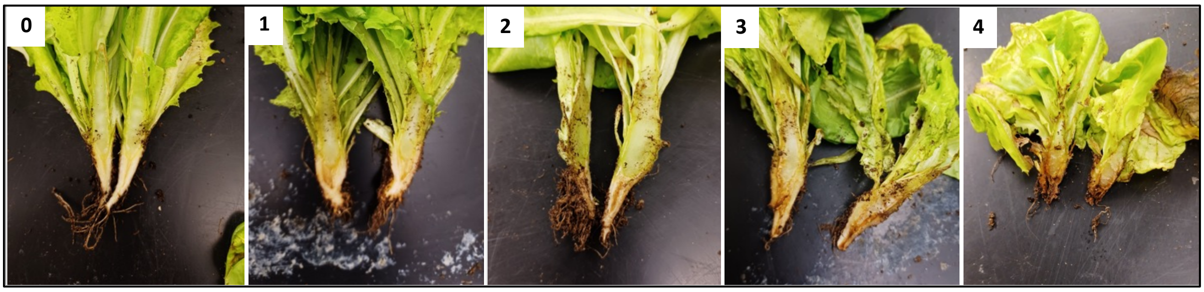


Figure X. Lettuce phenotyping information and disease scales: a) standard reactions of lettuce differential cultivars vs to Fola races (ISF Report, 2022); exemplar Fola disease symptoms for b) wilt and c) vascular browning disease scales.
